# Interplay between acetylation and ubiquitination of imitation switch chromatin remodeler Isw1 confers multidrug resistance in *Cryptococcus neoformans*

**DOI:** 10.1101/2022.12.30.522307

**Authors:** Yang Meng, Zhuoran Li, Tianhang Jiang, Tianshu Sun, Yanjian Li, Xindi Gao, Hailong Li, Chenhao Suo, Chao Li, Sheng Yang, Tian Lan, Guojian Liao, Tong-Bao Liu, Ping Wang, Chen Ding

## Abstract

*Cryptococcus neoformans* poses a threat to human health, but anticryptococcal therapy is hampered by the emergence of drug resistance, whose underlying mechanisms remain poorly understood. Herein, we discovered that Isw1, an imitation switch chromatin remodeling ATPase, functions as a master modulator of genes responsible for multidrug resistance in *C. neoformans*. Cells with the disrupted *ISW1* gene exhibited profound resistance to multiple antifungal drugs. Mass spectrometry analysis revealed that Isw1 is both acetylated and ubiquitinated, suggesting that an interplay between these two modification events exists to govern Isw1 function. Mutagenesis studies of acetylation and ubiquitination sites revealed that the acetylation status of Isw1^K97^ coordinates with its ubiquitination processes at Isw1^K113^ and Isw1^K441^ through modulating the interaction between Isw1 and Cdc4, an E3 ligase. Additionally, clinical isolates of *C. neoformans* overexpressing the degradation-resistant *ISW1*^*K97Q*^ allele showed impaired drug-resistant phenotypes. Collectively, our studies revealed a sophisticated acetylation-Isw1-ubiquitination regulation axis that controls multidrug resistance in *C. neoformans*.

## Introduction

Emerging and re-emerging fungal pathogens are one of the primary causes of infectious diseases resulting in mortalities in humans and animals (Brown et al., 2012; Denning and Bromley, 2015; Fisher et al., 2012; Vos et al., 2012). It is estimated that approximately 300 million world’s population is infected with fungal pathogens, subsequently leading to 1.6 million deaths annually (2017). Additionally, fungal infection leads to substantial population declines in animals and amphibians, even threatening their extinction (de Bekker et al., 2015; Fites et al., 2013; Martel et al., 2014; Voyles et al., 2009). Intriguingly, certain pathogenic fungi are linked with cancer development (Dohlman et al., 2022; Narunsky-Haziza et al., 2022). The World Health Organization has recently published the first-ever list of fungi posing threats to human health that lists *Cryptococcus neoformans* as one of four high-priority infection agents (2022). *Cryptococcus* species, which cause meningoencephalitis and acute pulmonary infections, are responsible for 15% of the global HIV/AIDS-related fatalities (Erwig and Gow, 2016; Kronstad et al., 2012), totalling an estimated 220,000 deaths per year (Erwig and Gow, 2016; Idnurm et al., 2005; Kronstad et al., 2012; Rajasingham et al., 2017). Recently, it has been reported that *Cryptococcus* is also involved in secondary infections in people with COVID-19 (Khatib et al., 2021; Woldie et al., 2020).

Cryptococcosis has a 100% death rate if patients are left untreated, but treatment remains challenging because the number of available antifungal medications is limited. Azoles and amphotericin B (Amp B) constitute the primary treatment options, with Amp B often in combination of 5-fluorocytosine (5-FC) (Molloy et al., 2018). In light of the treatment limitation and risks, as well as the high costs associated with developing new antifungal treatments, the US FDA has categorized anti-cryptococcus therapies as “orphan drugs,” granting regulatory support by reducing the requirements for clinical studies (Denning and Bromley, 2015). Nevertheless, resistance to anticryptococcal drugs occurs rapidly, outpacing the development of new therapeutic options.

The mechanisms by which antifungal resistance occurs can be classified as either mutation in drug targets or epigenetic phenomena (Billmyre et al., 2020; Li et al., 2020; van der Linden et al., 2011). Point mutations in binding domains or regions of the drug target proteins often hinder their interactions with drugs (Billmyre et al., 2020; Kwon-Chung and Chang, 2012; Li et al., 2020; Priest et al., 2022; van der Linden et al., 2011) resulting in resistance, such as in the case of lanosterol demethylase, encoded by the *ERG11* gene, in *Candida* and *Cryptococcus* species (Bosco-Borgeat et al., 2016; Sionov et al., 2012; Spruijtenburg et al., 2022). Additionally, pathogenic fungi display a disparate set of mutations that counteract various antifungal drugs. Defects in DNA mismatch repairs resulting in high gene mutation rates and, thereby, 5-FC resistance was identified in *C. deuterogattii*, a species distinct but close to *C. neoformans* (Billmyre et al., 2020). Mutations in genes encoding the cytosine permease (Fcy2), uracil phosphoribosyl transferase (Fur1), and UDP-glucuronic acid decarboxylase (Uxs1) conferring 5-FC resistance were also identified in multiple clinical cryptococcal isolates (Chang et al., 2021; Florent et al., 2009).

In comparison, nondrug target-induced resistance often refers to the association between mutations and altered gene expressions in regulatable components of sterol biosynthesis and efflux drug pumps (Coste et al., 2004; Dunkel et al., 2008; Morschhauser et al., 2007; Silver et al., 2004). Studies of *C. albicans* transcription factor Tac1 illustrate that gain-of-function mutations or alterations in gene copy numbers regulate efflux pump genes (Coste et al., 2004). Gain-of-function mutations were also identified in *C. albicans* transcription factors Upc2 and Mrr1, which activate *ERG11* gene expression and drug pumps, respectively (Dunkel et al., 2008; Silver et al., 2004). A cryptococcal Upc2 homolog was identified in *C. neoformans* to be involved in regulating steroid biosynthesis, but its role in drug resistance was not clear (Kim et al., 2010). Other proteins capable of modulating efflux pump gene transcription in *Cryptococcus* remain unknown.

Recent evidence indicates that epiregulation by protein posttranslational modification (PPTM) mediates drug resistance in pathogenic fungi (Calo et al., 2014; Robbins et al., 2012). The acetylation of the heat-shock protein Hsp90 mediates antifungal drug response in *Saccharomyces cerevisiae* and *C. albicans* by blocking the interaction between Hsp90 and calcineurin (Robbins et al., 2012). Despite this observation, the functions of substantially acetylated proteins in antifungal drug resistance remain unknown (Li et al., 2019). Other important PPTMs, such as ubiquitination, remain uninvestigated in human fungal pathogens.

In this study, we identified a conserved chromatin remodeler, Isw1, from *C. neoformans*, and we demonstrated its critical function in modulating gene expression responsible for multidrug resistance, as the *isw1* null mutant is resistant to azoles, 5-FC, and 5-fluorouracil (5-FU). We further demonstrated that Isw1 typifies a PPTM interplay between acetylation and ubiquitination in regulating an Isw1 ubiquitin-mediated proteasome axis in response to antifungal exposure. Dissection of PPTM sites on Isw1 revealed an essential reciprocal function of Isw1^K97^ acetylation in modulating Isw1’s binding to Cdc4, which initiates a ubiquitination-proteosome degradation process. Finally, we showed that the PPTM interplay mechanism occurs horizontally in clinical strains of *C. neoformans*.

## Results

### Cryptococcal Isw1 plays an indispensable role in modulating drug-resistance genes

We have found that a homolog of imitation switch (ISWI)-class ATPase subunit, Isw1, is a critical modulator of multidrug resistance in *C. neoformans*. We have generated the *isw1Δ* mutant and complemented the mutant with the wild-type *ISW1* allele (**Figure 1-figure supplement 1a-1c)**. The *isw1Δ* mutant exhibited normal phenotypes, similar to the wild-type strain, and was dispensable in the virulence of a murine infection model **(Figure 1-figure supplement 1d)**; however, it exhibited profound resistance to azole compounds, including fluconazole (FLC), ketoconazole (KTC), 5-FC, and 5-FU, but not to the polyene compound Amp B **(Figure 1a)**.

**Figure 1.**
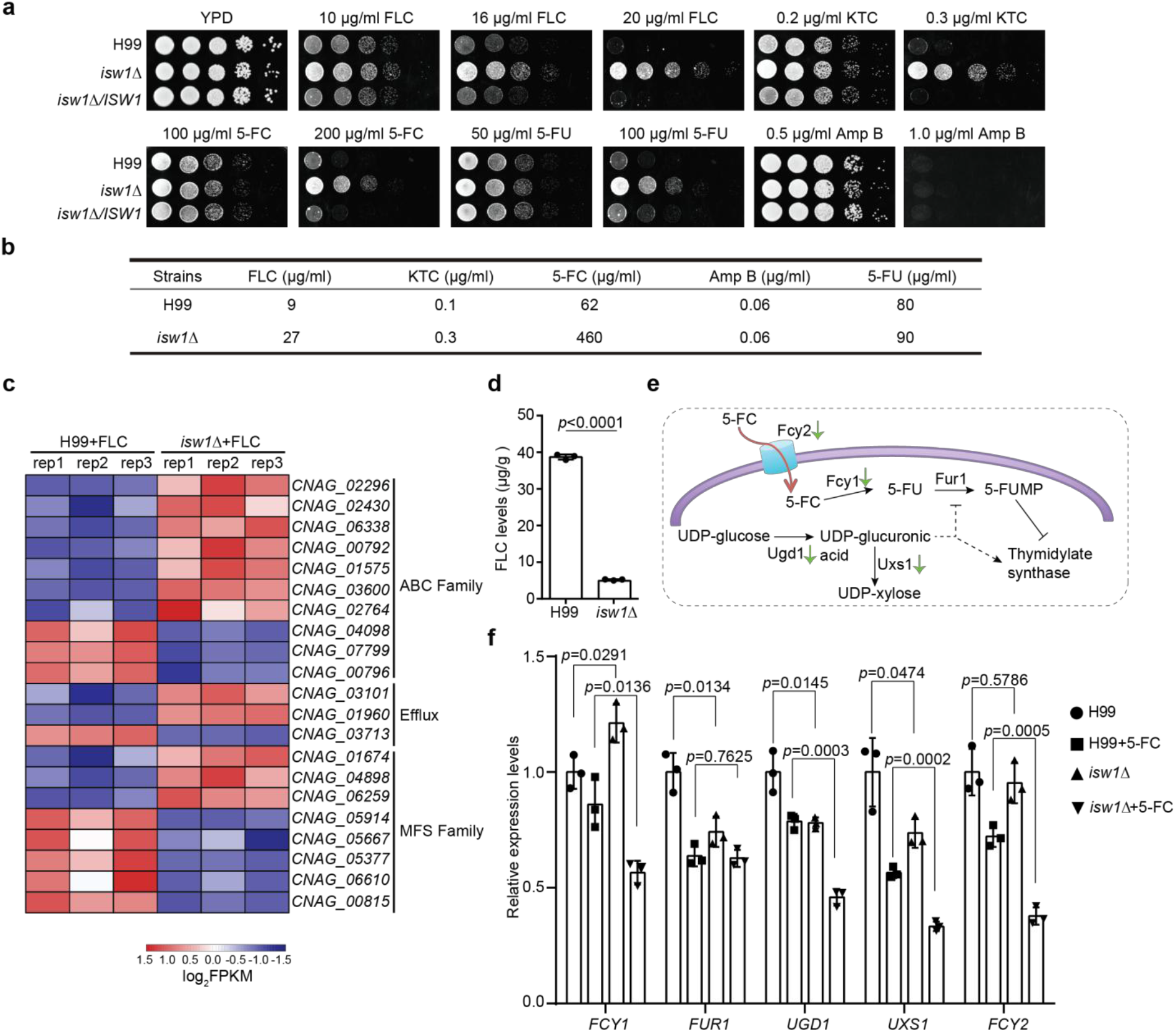
Isw1 represses the expression of drug-resistance genes. a. Spotting assays of *ISW1* mutant strains. Wild-type H99, *isw1Δ*, and *ISW1* complementation strains were spotted onto YPD agar supplemented with indicated concentrations of antifungal agents. Plates were incubated at 30°C for 3 days. b. Minimum inhibitory concentration (MIC) tests. The H99 and *isw1Δ* (*n*=3 each) strains were tested to determine the MICs of several antifungal agents. c. Transcriptome analysis of *isw1Δ*. Samples of RNA were isolated from H99 and *isw1Δ* cells (*n*=3 each) supplemented with 10 μg/ml fluconazole (FLC). Transcriptome analysis was performed, and a heat map of the expressions of drug-resistance genes was generated. d. Intracellular concentration of FLC. H99 and *isw1Δ* cells (*n*=3 each) were incubated in the presence of 40 μg/ml FLC at 30°C for 5 hours. Cells were then washed and weighed, and the intracellular FLC was quantified using high-performance liquid chromatography. A two-tailed unpaired *t*-test was used. Data are expressed as mean ± SD. e. Scheme of the mechanism for 5-fluorocytosine (5-FC) and 5-fluorouracil (5-FU) resistance. The red arrow indicates the entry of 5-FC via Fcy2. Green arrows indicate the downregulation of gene expression in response to 5-FC in the *isw1Δ* strain. f. Analyses of 5-FC resistance genes using qRT-PCR. Indicated strains (*n*=3 each) were grown with or without 400 μg/ml 5-FC, then qRT-PCR was performed to determine gene expressions of *FCY1, FUR1, UGD1, UXS1*, and *FCY2*. Two-tailed unpaired *t* tests were used. Data are expressed as mean ± SD. **Figure 1-figure supplement 1. *ISW1* is not required for fungal virulence. Figure 1-figure supplement 2. *itc1Δ* is resistant to azoles and 5-fluorocytosine. Figure 1-source data 1** **Figure 1-figure supplement 1-source data 1 Figure 1-figure supplement 2-source data 1**

Since the conserved ISWI complex often consists of two protein components, Isw1 and Itc1 (Sugiyama and Nikawa, 2001), we identified an Itc1 homolog from *C. neoformans* and generated the corresponding *itc1Δ* mutant **(Figure 1-figure supplement 2a and 2b)**. We found that the *itc1Δ* strain displayed a drug-resistant phenotype, similar to the *isw1Δ* strain, suggesting that Isw1 and Itc1 function together to modulate drug resistance **(Figure 1-figure supplement 2b)**. Both minimum inhibitory concentration (MIC) and agar spotting assays were consistent in showing the elevated MICs to all four antifungal compounds in the *isw1Δ* mutant strain **(Figure 1b)**. To examine whether Isw1-regulated multidrug resistance occurs at the transcription level, transcriptome analysis of the wild-type and *isw1Δ* mutant strains treated with FLC was performed **(Table S1)**. The expression of genes important in drug resistance (Denning and Bromley, 2015), including seven genes encoding ATP-binding cassette (ABC) transporters, two genes encoding efflux proteins, and three genes encoding major facilitator superfamily (MFS) transporters, were significantly increased in the *isw1Δ* mutant **(Figure 1c)**. To test if this increase affects drug uptake, we quantified intracellular FLC levels using high-performance liquid chromatography that showed a significant reduction in FLC in the *isw1Δ* mutant, in contrast to the wild-type strain **(Figure 1d)**.

To further decipher the 5-FC-resistance mechanisms of *isw1Δ*, we examined 5-FC resistance pathways. As previously elucidated, *C. neoformans* employs two molecular processes for resistance to 5-FC and 5-FU (Billmyre et al., 2020; Loyse et al., 2013). In one, the purine-cytosine permease Fcy2 imports the prodrug 5-FC, which is then converted to toxic 5-FU by the purine-cytosine permease Fcy1 **(Figure 1e)**. On the other, the UDP-glucose dehydrogenase Ugd1 and the UDP-glucuronate decarboxylase Uxs1 participate in UDP-glucose metabolism, providing important functions in detoxifying 5-FU **(Figure 1e)**. Therefore, gene expression alterations in *FCY1, FCY2, FUR1, UGD1*, and *UXS1* were assayed using the qRT-PCR method **(Figure 1f)**. The data showed that the expression of four genes, including *FCY1, FCY2, UGD1*, and *UXS1*, was significantly reduced in *isw1Δ* when treated with 5-FC, suggesting a reduction in 5-FC uptake and conversion to toxic 5-FU **(Figure 1f)**. These data suggested that Isw1 is a master transcriptional regulator of drug-resistance genes in *C. neoformans*.

### Isw1 undergoes protein degradation in the presence of azoles and 5-FC

Because Isw1 governs the expression of multiple genes required for drug resistance and *ISW1* gene expression was not reduced in the presence of antifungal drugs **(Figure 1-figure supplement 2c)**, we examined changes in protein stability as a response to antifungal agents. Using cycloheximide to inhibit protein synthesis, Isw1-Flag fusion protein stability was found to decrease gradually in concentration- and time-dependent manners upon exposure to FLC **(Figures 2a and 2b)**. Specifically, a reduction of 50% was observed 30 minutes after FLC exposure. Similarly, 5-FC exposure also reduces Isw1-Flag levels **(Figures 2c and 2d)**. Collectively, these results demonstrated that *C. neoformans* actively reduces Isw1 protein levels through protein degradation than transcription to manage toxicity overloads from antifungals.

**Figure 2.**
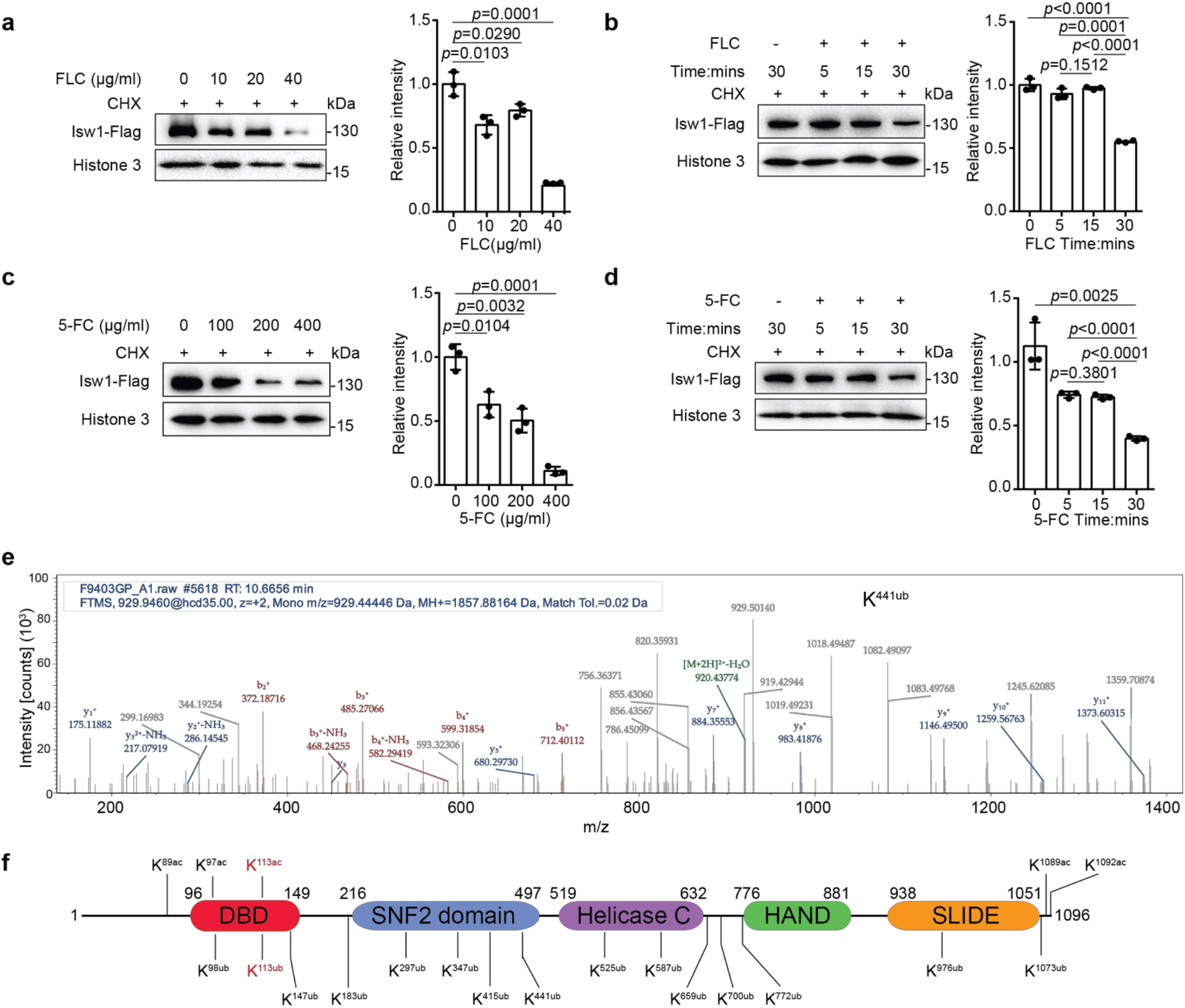
Isw1 is an acetylated and ubiquitinated protein. a. Immunoblotting analysis. The *ISW1-FLAG* strain was preincubated with 200 μM cycloheximide for 1 hour followed by exposure to various concentrations of fluconazole (FLC) for 0.5 hours. Anti-Flag and anti-histone 3 antibodies were used. Three biological replicates were performed, and results were used for quantification. Two-tailed unpaired *t*-tests were used. Data are expressed as mean ± SD. b. Immunoblotting analysis. The *ISW1-FLAG* strain was preincubated with 200 μM cycloheximide for 1 hour followed by exposure to 40 μg/ml FLC for 5, 15 or 30 minutes. Cells not exposed to FLC but held for 30 minutes were used as a negative control. Anti-Flag and anti-histone 3 antibodies were used. Three biological replicates were performed, and results were used for quantification. Two-tailed unpaired *t* tests were used. Data are expressed as mean ± SD. c. Immunoblotting analysis. Testing and data treatment were exactly as described for Figure 2a with the exception that 5-fluorocytosine (5-FC) was used as the antifungal agent. d. Immunoblotting analysis. Testing and data treatment were exactly as described in Figure 2b, except that 5-FC was used as the antifungal agent. e. Ubiquitin analysis of Isw1 via mass spectrometry. The Isw1-Flag proteins were pulled down and analyzed for ubiquitination. Results for Isw1^K441Ub^ are shown. f. Schematic of Isw1 showing acetylation (Li et al., 2019) and ubiquitination sites. **Figure 2-source data 1**

We then hypothesized that Isw1 degradation might be via a ubiquitin-proteasome pathway in response to antifungal drugs. To test this, the Isw1-Flag protein was immunoprecipitated and then analyzed using mass spectrometry to identify putative ubiquitination PPTM sites. Consistent with our hypothesis, Isw1 is ubiquitinated **(Figure 2e)** at fifteen sites **(Figure 2f)**. These results indicated that the ubiquitination machinery of Isw1 is actively initiated during drug exposure, and this, in turn, decreases Isw1 protein levels and hinders Isw1 transcription repression of genes for drug resistance. The finding of Isw1 subject to ubiquitination and acetylation regulation (Li et al., 2019) also suggests that there exists an interplay network simultaneously controlling Isw1 stability in response to antifungal drugs. We then set forth to address the following two questions: 1) whether there is a PPTM interplay between acetylation and ubiquitination in Isw1; and 2) how Isw1 utilizes this interplay regulates responses to antifungals.

### Acetylation of Isw1^K97^ (Isw1^K97ac^) is essential for protein stability

To dissect the interplay between acetylation and ubiquitination in Isw1, we examined the role of acetylation in modulating Isw1 function by determining acetylation levels responding to antifungal drugs. The presence of antifungal agents strongly repressed acetylation levels, in contrast to deacetylation inhibitors trichostatin A (TSA) and nicotinamide (NAM), which increase acetylation levels **(Figures 3a and 3b)**. These data suggested a positively regulated deacetylation process in Isw1 in response to antifungal drugs. To more closely decipher this regulation mechanism, three acetylation sites, K89, K97, and K113, located to the DNA binding domain were mutated to arginine (R) to mimic a deacetylated status or to glutamine (Q) to mimic a fully acetylated Isw1 **(Figure 2f)**. Gene copy numbers and transcription levels were confirmed to be equivalent to those of the wild-type strain **(Figure 3-figure supplement 1a and 1b)**. Triple-, double-, and single-mutated strains were generated, and their drug resistance phenotypes were compared. Of the triple-mutated strains, cells with three R mutations demonstrated drug-resistant growth phenotypes that were similar to those of the *isw1Δ* strain, whereas those with three Q mutations showed wild-type growth in the presence of antifungal drugs **(Figure 3c)**. Of the double-mutated strains, *ISW1*^*K89R, K97R*^, and *ISW1*^*K97R, K113R*^ strains showed resistance to antifungal drugs, with *ISW1*^*K97R, K113R*^ less so than *ISW1*^*K89R, K97R*^ or the *isw1Δ* strain **(Figure 3-figure supplement 1c)**. These data suggested that the acetylation status of Isw1^K97^ is important in conferring drug resistance. Of the strains that have single-R mutation, the *ISW1*^*K97R*^ strain showed robust resistance to antifungal drugs, mimicking the *isw1Δ* strain **(Figure 3d)**. Interestingly, the *ISW1*^*K97Q*^ strain showed no drug resistance **(Figure 3-figure supplement 1d)**. Collectively, these data strongly demonstrated that the acetylation status of Isw1^K97^ plays a critical role in regulating Isw1 protein stability and function in response to antifungal drugs.

**Figure 3.**
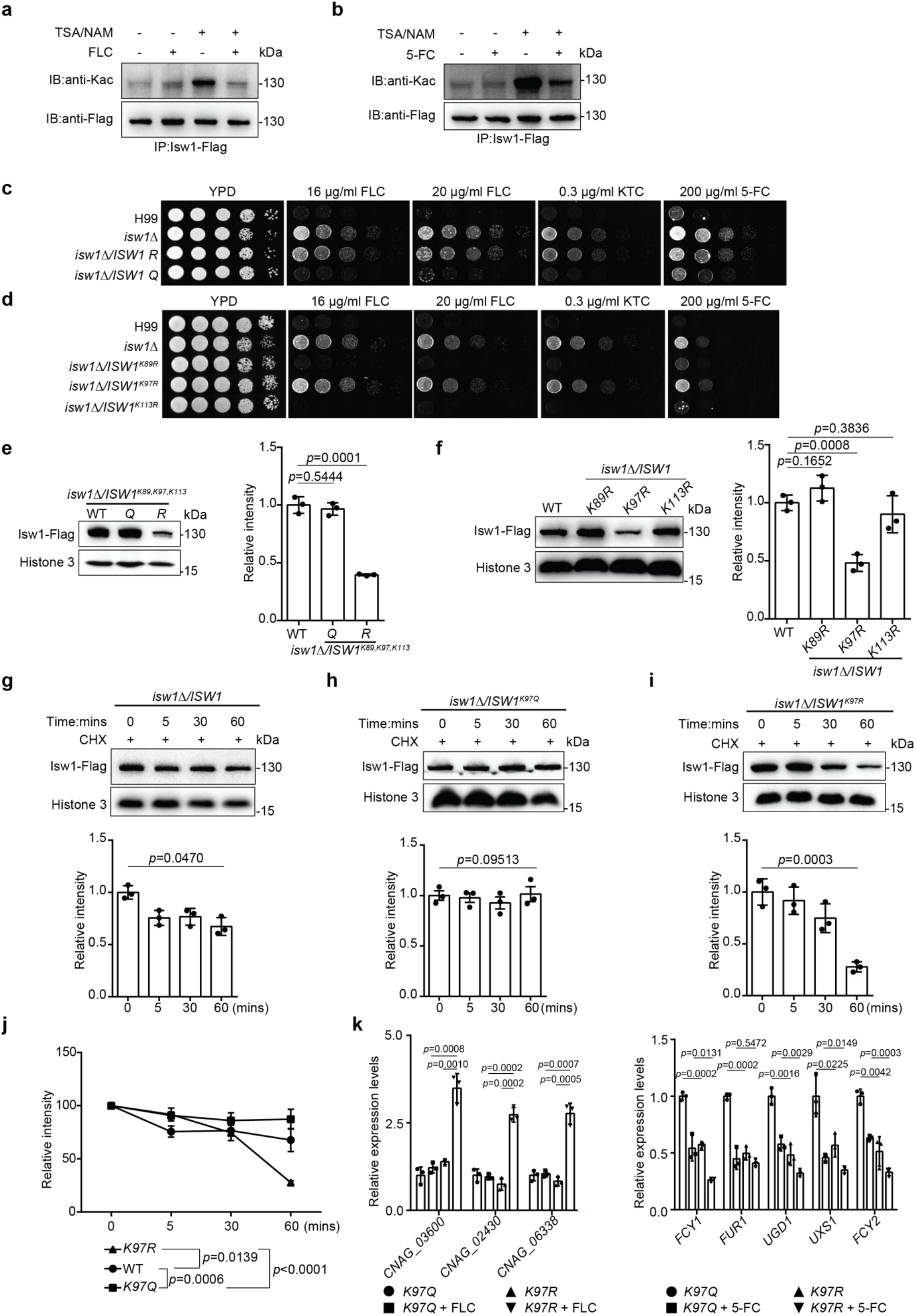
The acetylation status of Isw1^K97^ (Isw1^K97ac^) is essential in Isw1 protein stability. a. Acetylation analysis of Isw1. Cells were treated with 3 μM trichostatin A (TSA), 20 mM nicotinamide (NAM), and fluconazole (FLC). The Isw1-Flag proteins were pulled down, and immunoblotting assays were performed using anti-Kac and anti-Flag antibodies. b. Acetylation analysis of Isw1. Cells were treated with TSA, NAM, and 5-fluorocytosine (5-FC). The Isw1-Flag proteins were pulled down, and immunoblotting assays were performed using anti-Kac and anti-Flag antibodies. c. Spotting assays of *ISW1* mutants. The *ISW1*^*K89R, K97R, K113R*^ and *ISW1*^*K89Q, K97Q, K113Q*^ strains were tested for drug resistance. d. Spotting assays of *ISW1* mutants. The *ISW1*^*K89R*^, *ISW1*^*K97R*^, and *ISW1*^*K113R*^ strains were tested for drug resistance. e. Immunoblotting assays of *ISW1* mutants. The wild-type, *ISW1*^*K89R, K97R, K113R*^, and *ISW1*^*K89Q, K97Q, K113Q*^ strains were tested for Isw1 levels. Three biological replicates were performed, and results were used for quantification. Two-tailed unpaired *t*-tests were used. Data are expressed as mean ± SD. f. Immunoblotting assays of *ISW1* mutants. The wild-type, *ISW1*^*K89R*^, *ISW1*^*K97R*^, and *ISW1*^*K113R*^ strains were tested for Isw1 levels. Three biological replicates were performed, and results were used for quantification. Two-tailed unpaired *t*-tests were used. Data are expressed as mean ± SD. g. Immunoblotting assay of Isw1. The wild-type strain was preincubated with 200 μM cycloheximide for 1 hour. Proteins were isolated at indicated time points. Three biological replicates of immunoblotting were performed, and results were used for quantification. One-way ANOVA was used. h. Immunoblotting assay of Isw1^K97Q^. The analysis was performed as described in Figure 3g. i. Immunoblotting assay of Isw1^K97R^. The analysis was performed as described in Figure 3g. j. Comparisons of assay results to determine Isw1 stability. The relative intensities from the results shown in Figures 3g, 3h, and 3i were plotted. Two-way ANOVA was used. Data are expressed as mean ± SD. k. Analyses of drug resistance genes using qRT-PCR. Samples of RNA (*n*=3) were isolated from *ISW1*^*K97Q*^ and *ISW1*^*K97R*^ treated with FLC or 5-FC. Representative drug resistance genes were quantified using qRT-PCR. Two-tailed unpaired *t*-tests were used. Data are expressed as mean ± SD. **Figure 3-figure supplement 1. Screening important acetylation sites of Isw1. Figure 3-source data 1.** **Figure 3-figure supplement 1-source data 1**.

To further investigate how Isw1 degradation correlates with drug-resistance of *C. neoformans*, we tested how Isw1^K97^ acetylation affects its degradation using the immunoblotting method, and the results showed that triple-R mutation results in a significant reduction in levels of Isw1-Flag. Meanwhile, triple-Q mutation resulted in Isw1-Flag levels comparable to those of the wild-type strains **(Figure 3e)**. Similarly, the single-mutated *ISW1*^*K97R*^ strain showed an impairment level of Isw1-Flag **(Figure 3f)**. In contrast, no changes were observed for Isw1^K97Q^ **(Figure 3-figure supplement 1e)**. Moreover, protein levels of wild-type Isw1 and mutated Isw1^K97R^ gradually diminished over time. While those of Isw1^K97Q^ remained constant **(Figures 3g, 3h, and 3i)**, a faster degradation was observed for Isw1^K97R^ **(Figure 3j)**. Therefore, Isw1^K97^ is an essential regulation site responsible for Isw1 stability; that is, acetylation at K97 blocks the degradation of Isw1, and deacetylation at K97 facilitates and accelerates the degradation of Isw1. Finally, analysis of Isw1 target gene expression in Isw1^K97^ mutation strains demonstrated significantly increased expression of transporter genes and decreased expression of 5-FC resistance genes **(Figure 3k)**. These findings were consistent with the results of transcriptome and qRT-PCR analyses in the *isw1Δ* strain **(Figures 1c and 1f)**.

### The interplay between acetylation and ubiquitination governs Isw1 degradation

As proteins undergo degradation via autophagy and proteasomal pathways, we employed autophagy inhibitor rapamycin and proteasome inhibitor MG132 to investigate Isw1 degradation. Immunoblotting results showed that both rapamycin and MG132 induce Isw1-Flag levels, with the effect of MG132 being stronger than rapamycin **(Figure 4a)**. These findings suggested that both degradation pathways are utilized in Isw1-Flag degradation, but the ubiquitin-mediated proteasomal process has a more predominant role. Additionally, we tested whether antifungal agents could induce protein degradation when the proteasome is blocked. Cells treated with MG132 and FLC or 5-FC yielded slightly different results. Isw1-Flag protein levels were unaffected in cells treated with FLC, but the levels were reduced with 5-FC **(Figures 4b and 4c)**.

Given that K97 deacetylation could trigger hyper-ubiquitination of Isw1, we analyzed Isw1 ubiquitination sites and their regulation mechanisms by K97 acetylation levels and found that MG132 treatment results in a more robust increase in Isw1^K97R^-Flag levels **(Figure 4d)**. We further performed ubiquitination site mutations in the genetic background of *ISW1*^*K97R*^ and found that, of the six ubiquitination sites **(Figure 4-figure supplement 1a and 1b)**, five failed to affect drug-resistant growth phenotypes of the *ISW1*^*K97R*^ mutant **(Figure 4-figure supplement 1c)**. Only *ISW1*^*K113R*^ and *ISW1*^*K441R*^ mutations exhibited reduced drug-resistant growth **(Figure 4e)**, indicating that they affect drug resistance by modulating Isw1 protein stability. The immunoblotting analysis further showed that, while all ubiquitination mutants exhibiting increased Isw1-Flag levels **(Figures 4f and 4g)**, the elevation was most pronounced in Isw1^K97R, K113R^ (5.6-fold) and Isw1^K97R, K441R^ (14.5-fold) **(Figures 4f and 4g)**. Interestingly, the K113 site may undergo acetylation or ubiquitination modifications, whereas the K441 site undergoes only ubiquitination. Collectively, these results showed that Isw1^K113^ and Isw1^K441^ provide a predominant role in regulating the ubiquitin-proteasome process of Isw1 and that acetylation at Isw1^K97^ has a broad role in controlling the ubiquitination process at those sites.

**Figure 4.**
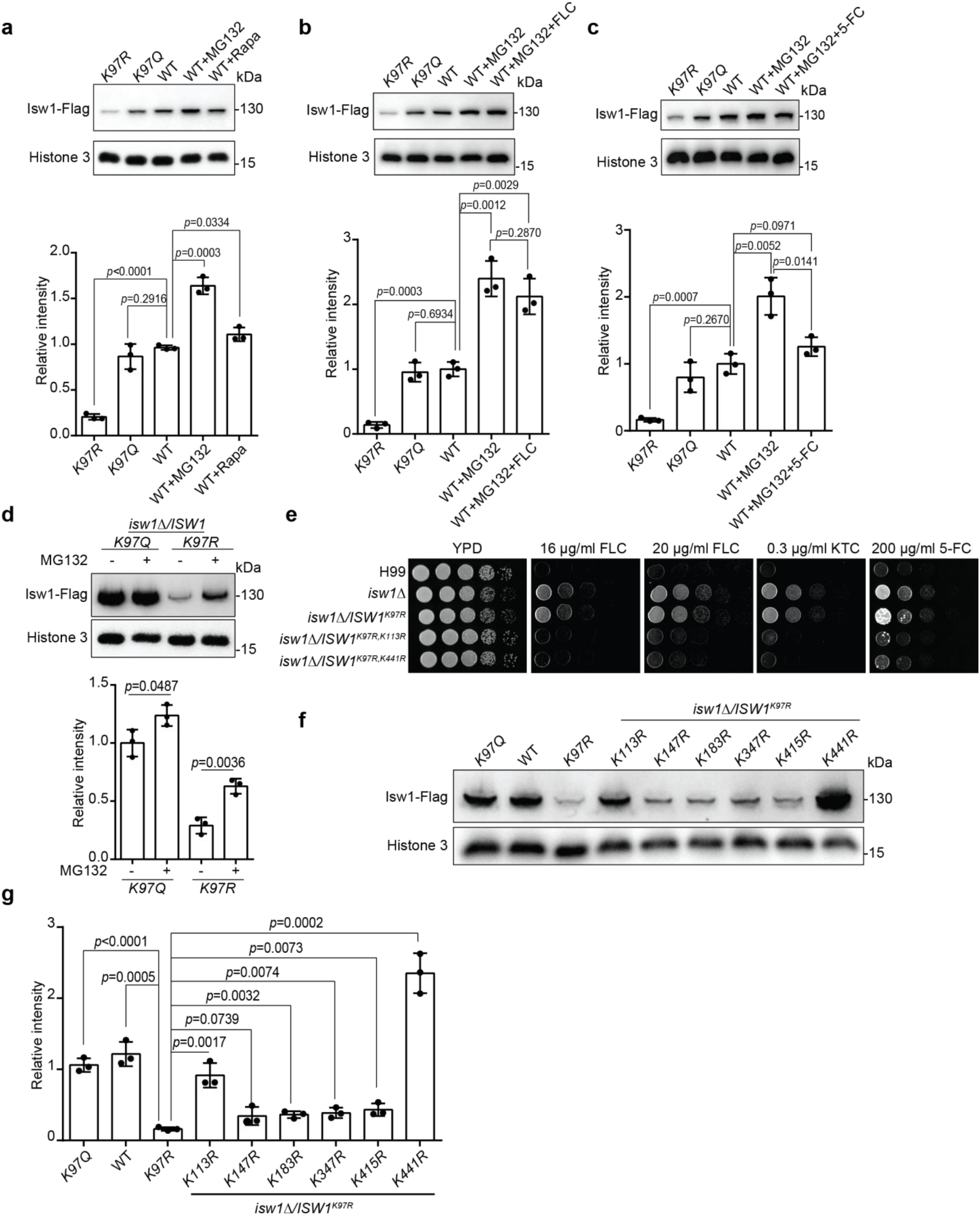
Isw1^K97ac^ is critical for Isw1 ubiquitin-proteasome degradation. a. Immunoblotting assays of Isw1-Flag. Proteins Isw1^K97Q^, Isw1^K97R^, and Isw1 were tested, where the wild-type stain was incubated with 200 μM MG132 and 5 nM rapamycin for 10 hours. Three biological replicates of the assays were performed, and results were used for quantification. Two-tailed unpaired *t* tests were used. Data are expressed as mean ± SD. b. Immunoblotting assays of Isw1-Flag. Testing and data treatment were exactly as described in Figure 4a, except that the wild-type sample was treated with 200 μM of MG132 and 40 μg/ml fluconazole (FLC). c. Immunoblotting assays of Isw1-Flag. Testing and data treatment were exactly as described in Figure 4a, except that the wild-type sample was treated with 200 μM MG132 and 400 μg/ml 5-fluorocytosine (5-FC). d. Immunoblotting assays of Isw1^K97Q^ and Isw1^K97R^. Proteins were either treated with MG132 or not before testing. Three biological replicates of the assays were performed, and results were used for quantification. Two-tailed unpaired *t-*tests were used. Data are expressed as mean ± SD. e. Spotting assays of *ISW1* acetylation and ubiquitination mutants. Indicated strains were spotted onto YPD agar either supplemented with an antifungal agent or left blank. f. Immunoblotting assays of *ISW1* acetylation and ubiquitination mutants. Protein samples were isolated from the indicated *ISW1* mutants. Immunoblotting assays were performed. g. Quantification of immunoblotting results. The immunoblotting assays described for Figure 3f were performed using three independent samples, and the results were used for quantification. Two-tailed unpaired *t-*tests were used. Data are expressed as mean ± SD. **Figure 4-figure supplement 1. Screening important ubiquitination sites of Isw1. Figure 4-source data 1.** **Figure 4-figure supplement 1-source data 1**.

### The acetylation status of Isw1^K97^ modulates the binding of an E3 ligase to Isw1

To analyze the molecular mechanism by which Isw1^K97^ regulates the ubiquitin-proteasome process, we generated nine knockout mutants of E3 ligase encoding genes in the genetic background of *ISW1*^*K97R*^ and identified that Cdc4 is an E3 ligase for Isw1 **(Figure 5-figure supplement 1 and Figure 5a)**. We found that the *ISW1*^*K97R*^*/cdc4Δ* strain becomes sensitive to antifungal agents **(Figure 5a)**. Immunoblotting showed a strong elevation in Isw1^K97R^-Flag levels in the *ISW1*^*K97R*^*/cdc4Δ* strain **(Figure 5b)**, in contrast to the *ISW1*^*K97R*^*/fwd1Δ* strain **(Figure 5c)**, suggesting a potential interaction between Cdc4 and Isw1. We then performed a co-immunoprecipitation (co-IP) assay and found that Cdc4-HA co-precipitates with Isw1-Flag **(Figure 5d)**.

**Figure 5.**
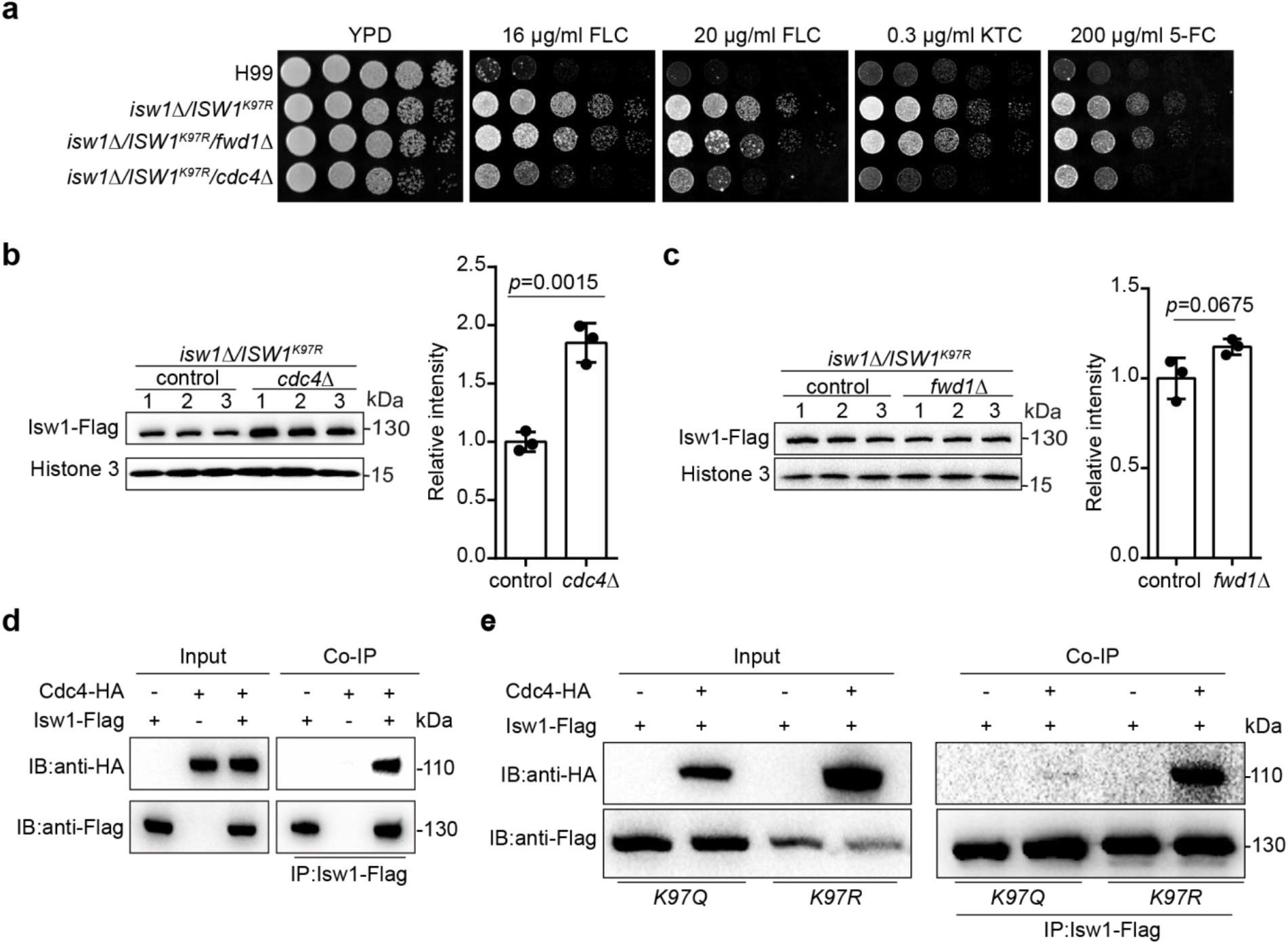
Isw^K97ac^ blocks the binding of Isw1 to the E3 ligase Cdc4. a. Spotting assays of E3 ligase mutants. Indicated strains were spotted onto YPD agar either supplemented with an antifungal agent or left blank. b. Immunoblotting assays of *cdc4Δ*. Protein samples were isolated from *isw1Δ /ISW1*^*K97R*^/*cdc4Δ* and its relevant control strains. Three independent samples were tested and quantified. Two-tailed unpaired *t-*tests were used. Data are expressed as mean ± SD. c. Immunoblotting assays of *fwd1Δ*. Protein samples were isolated from *isw1Δ /ISW1*^*K97R*^/*fwd1Δ* and its relevant control strains. Three independent samples were tested and quantified. Two-tailed unpaired *t-*tests were used. Data are expressed as mean ± SD. d. Protein co-immunoprecipitation (co-IP) of Cdc4 and Isw1. Protein samples were isolated from the strain expressing Cdc4-HA and Isw1-Flag, and co-IP was performed. e. Protein co-IP of Cdc4 and Isw1 K97 mutant proteins. Protein samples were isolated from the strain co-expressing Cdc4-HA and Isw1^K97R^-Flag and the strain co-expressing Cdc4-HA and Isw1^K97Q^-Flag. Co-IP was performed for each. **Figure 5-figure supplement 1. Identification of the E3 ligase for Isw1. Figure 5-source data 1**.

We also carried out co-IP to examine interactions between Cdc4-HA and Isw1^K97R^ and Cdc4-HA and Isw1^K97Q^, and the results showed an interaction between Cdc4 and Isw1^K97R^ but not between Cdc4 and Isw1^K97Q^, indicating the acetylation of Isw1^K97^ fully hinders its binding of the Cdc4 E3 ligase **(Figure 5e)**. These data provided convincing evidence that K97 acetylation is a key player in modulating ubiquitin-proteasome degradation of Isw1.

### The Isw1-proteasome regulation axis promotes drug resistance in clinical isolates

We have provided evidence demonstrating the critical role of Isw1 and Isw1 acetylation-ubiquitin-proteasome regulation axis in regulating *C. neoformans* drug resistance. To examine whether such a regulatory role has a broad application, we tested various clinical isolates of *C. neoformans*. While the majority was resistant to antifungal drug exposure, the CDLC120 isolate showed moderate sensitivity to all tested antifungal drugs **(Figure 6a)**. An integrative Flag-tag construct was then transformed into several clinical strains, allowing the expression of Isw1-Flag from the *ISW1* endogenous promoter. Expression analysis showed that transformants exhibiting multidrug resistance phenotypes have significantly reduced levels of Isw1-Flag **(Figure 6b)**. Importantly, the reduction was not at the transcription level **(Figure 6c)**. In contrast, the CDLC120 isolate showed increased Isw1-Flag levels. Similar to the PPTMs of Isw1 from the wild-type strain, samples of Isw1 from clinical isolates showed elevated acetylation levels in response to TSA and NAM **(Figure 6d)**. Moreover, clinical isolates treated with MG132 showed improved Isw1 stability **(Figure 6e)**. The clinical isolates were then examined to determine whether drug resistance results from changes in Isw1 protein levels following transformation with an integrative plasmid overexpressing *ISW1*^*K97Q*^. The results revealed decreased MICs for antifungal agents in these strains **(Figure 6f)**. Therefore, Isw1 acetylation-ubiquitin-proteasome regulation axis is a naturally occurring strategy for modulating multidrug resistance in clinical strains of *C. neoformans*.

**Figure 6.**
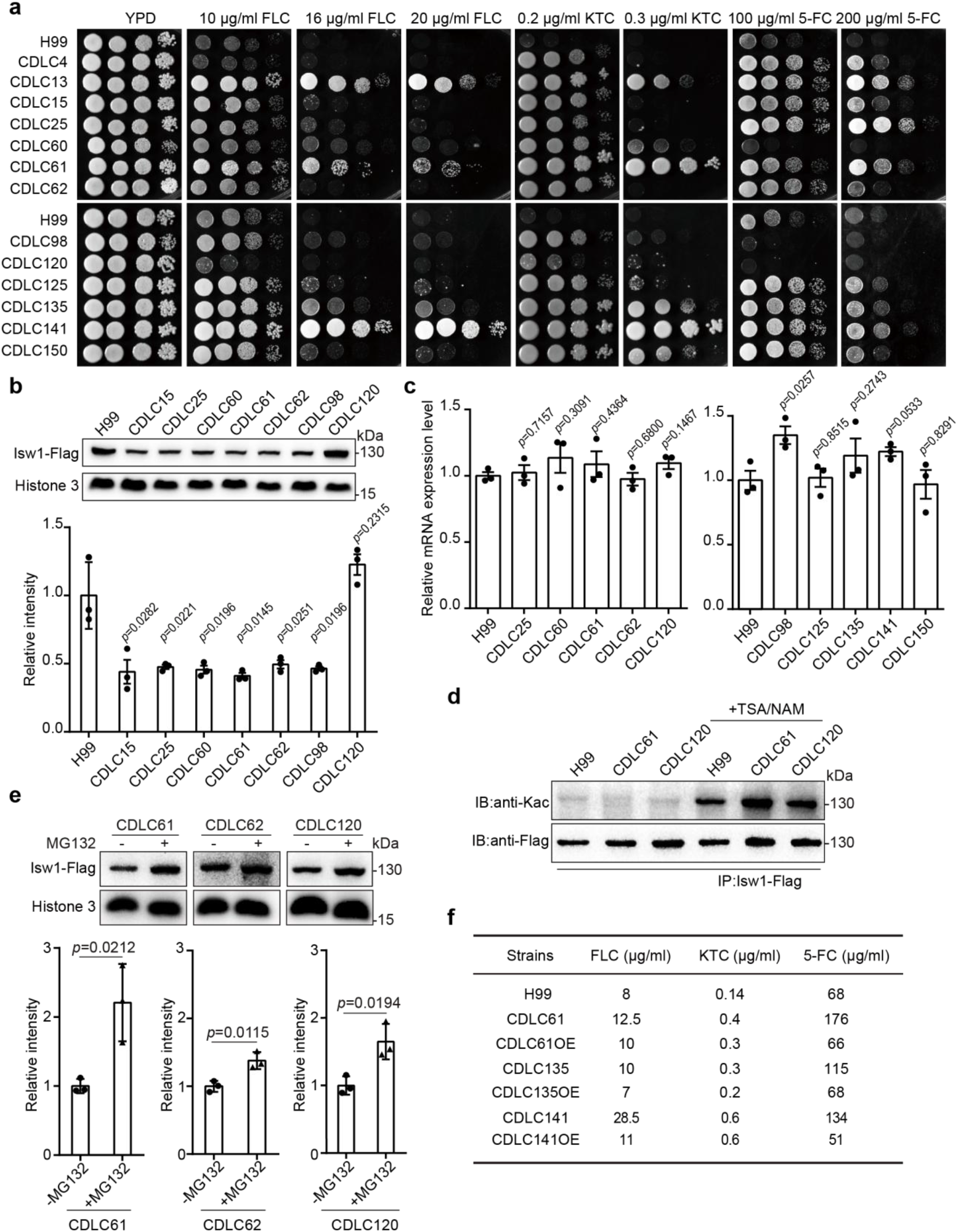
Clinical *C. neoformans* isolates show Isw1-mediated drug resistance phenotypes. a. Spotting assays of *C. neoformans* clinical strains. Clinical strains were spotted onto YPD agar either supplemented with an antifungal agent or left blank. Results after 3 days of incubation at 30°C are shown. b. Immunoblotting assays of Isw1-Flag from clinical isolates. Protein samples were isolated from H99 and clinical isolates expressing Isw1-Flag. Immunoblotting analyses were performed. Three independent repetitions were performed, and results were used for quantification. Two-tailed unpaired *t-*tests were used. Data are expressed as mean ± SD. c. Analyses of *ISW1* in clinical isolates using qRT-PCR. Samples of RNA were isolated from H99 and clinical strains, then qRT-PCR performed on each followed by quantification of *ISW1*. Three biological replicates were performed. Two-tailed unpaired *t-*tests were used. Data are expressed as mean ± SD. d. Immunoblotting analyses of Isw1 acetylation in clinical strains. Cells were treated with trichostatin A and nicotinamide, then Isw1-Flag was immunoprecipitated. Acetylation levels were examined using anti-Kac antibodies. e. Immunoblotting analyses of Isw1 levels under MG132 treatment. Cells were treated with MG132, then immunoblotting assays were performed on Isw1-Flag using anti-Flag antibodies. Three independent repetitions were performed, and results were used for quantification. Two-tailed unpaired *t-*tests were used. Data are expressed as mean ± SD. f. Minimum inhibitor concentration (MIC) analyses of Isw1-overexpressing clinical strains. Clinical strains harboring integrative overexpressing plasmid of *ISW1*^*K97Q*^ were tested for drug resistance, and MICs were determined. OE indicates strains with overexpressed *ISW1*^*K97Q*^. **Figure 6-source data 1**.

## Discussion

Fungi have developed sophisticated machinery to combat various stress inducers, and the rapid emergence of resistance to antifungal agents is one of the major factors in the failure of clinical therapies for fungal infections (Denning and Bromley, 2015). A typical tactic used by fungi to overcome antifungal toxicity is to utilize polymorphisms or mutations in drug targets or their regulatory components. Clinical polymorphisms were widely shown in drug targets, such as ergosterol biosynthesis and its transcription regulatory process (Billmyre et al., 2020; Denning and Bromley, 2015). However, unlike immediate intracellular responses, the accumulation of mutations or polymorphisms in drug resistance is a somewhat delayed process that frequently develops over a series of cell divisions. Acetylation and ubiquitination are critical modulators of protein activities or stabilities in fungi that enable rapid intracellular adaptations to environmental or chemical stressors (Li et al., 2019; Wu et al., 2021), but the underlying mechanisms are not clear. Recently, a study showed that the deactivation of a heat-shock Hsp90 client protein and its stability as a result of changes in protein acetylation impacts drug resistance in *C. albicans* (Robbins et al., 2012). Additional studies have demonstrated that deacetylation enzymes, notably Gcn5, control biofilm formation, morphology, and susceptibility to antifungal drugs in several fungi (O’Meara et al., 2010; Rashid et al., 2022; Yu et al., 2022). Despite this, knowledge of the molecular machinery of posttranslational modifications in modulating drug resistance remains not clear.

We demonstrated that the chromatin remodeler Isw1 is a master regulator of drug resistance in *C. neoformans*, and the acetylation-Isw1-ubiquitination axis is crucial in modulating the expression of multiple drug resistance genes. In *S. cerevisiae*, Isw1 is a key component of the ISWI complex capable of forming complexes with loc3, loc4, Itc1, and other proteins to modulate transcription initiation and elongation (Sugiyama and Nikawa, 2001; Tsukiyama et al., 1999). In *C. neoformans*, transcriptome analysis revealed 1275 genes, approximately 18.3% of the genome, that were significantly differentially expressed in the *isw1Δ* mutant treated with FLC. These alterations in gene expression activate the transcription of twelve drug pump genes, including ABC and MFS transporter genes, followed by a drastic reduction in intracellular FLC levels. The robust drug resistance phenotype (3-fold compared to the wild-type) of *isw1Δ* is the result of the simultaneous activation of various drug pumps, which then actively eliminate intracellular drug molecules. The expressions of genes required for resisting 5-FC and 5-FU were reduced when cells were treated with 5-FC. While Isw1 is a transcription activator for genes responsible for resistance to FLC, it also functions as a repressor in the presence of 5-FC, implying that transcriptional remodeling of Isw1 is necessary as the cell responds to disparate chemical stresses and that Isw1 engages distinct regulatory venues to overcome drug toxicity.

We also showed that the Isw1 protein and its acetylation level act reciprocally to govern fungal drug resistance **(Figure 7)**. This was confirmed by uncovering the interplay mechanism between acetylation and ubiquitination. The total acetylation levels of Isw1 were reduced when cells were treated with FLC or 5-FC, leading to the activation of Isw1 ubiquitination machinery. We identified that the K97 acetylation site functions as the essential regulating component of this interplay. We also found that K97 acetylation acts as a switch for ubiquitin conjugation proceeding proteasome-mediated degradation. When deacetylated, K97 triggers the activation of Isw1 degradation via the ubiquitin-proteasome process (acetylated K97 blocks the physical interaction with Cdc4). Compared to the *ISW1*^*K97R*^ strain, *ISW1*^*K97R*^/*cdc4Δ* was sensitive to antifungal agents; however, it showed moderately resistant growth in comparison to the wild-type strain. This data suggested that Isw1 could also be modulated by other proteins, such as another uncharacterized E3 ligase. Such hypothesis is supported by evidence from the comparison of Isw1 ubiquitination mutants with those of the *ISW1*^*K97R*^/*cdc4Δ* strain, and from that, the Isw1^K441R^ mutant had a 14.5-fold increase of Isw1 and a 2-fold increase in the *ISW1*^*K97R*^/*cdc4Δ* strain.

**Figure 7.**
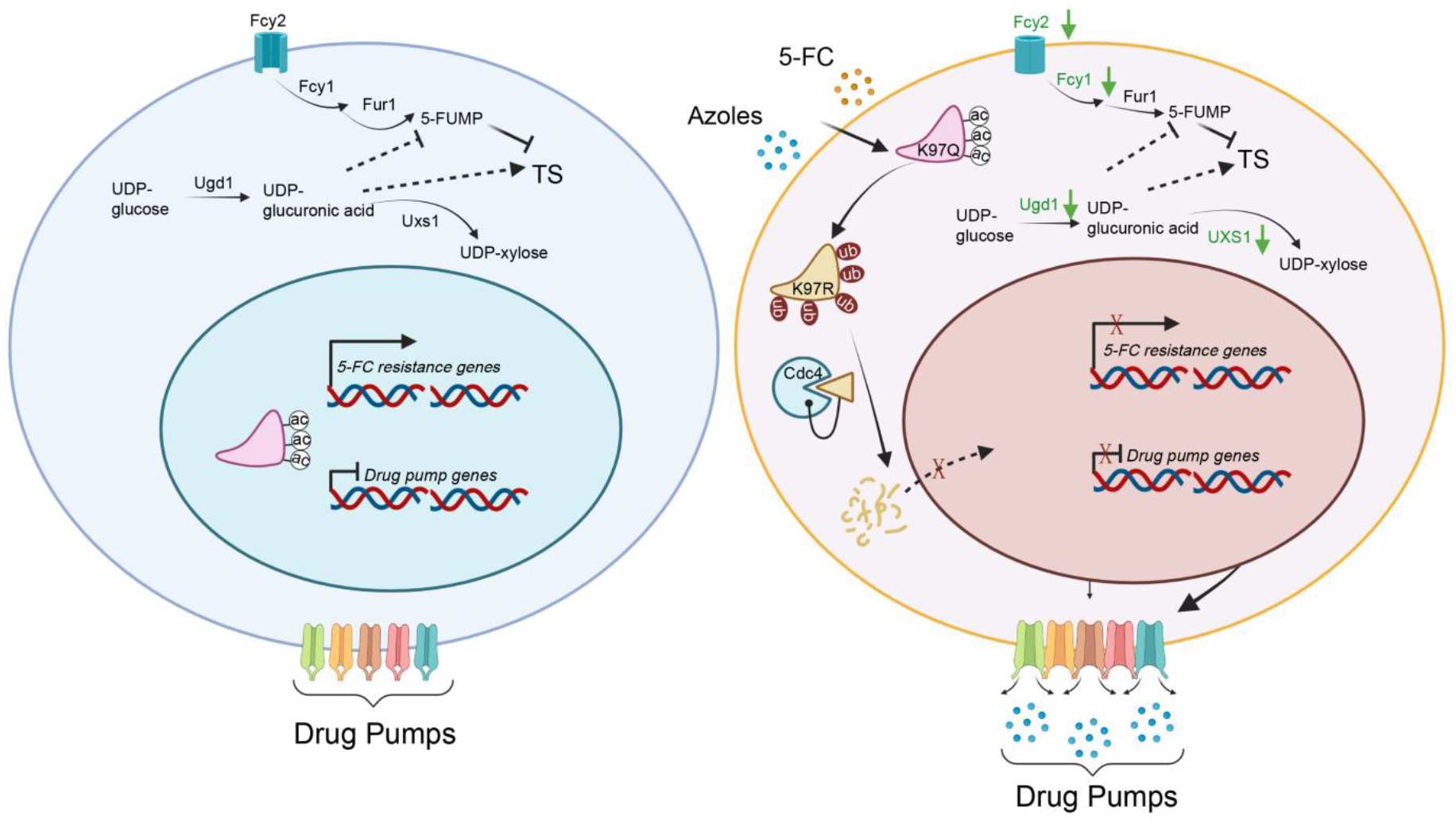
A model of the mechanism of Isw1 PPTM interaction in *C. neoformans* drug resistance. In a drug-free environment, acetylated Isw1 regulates 5-FC resistance gene expression and represses drug pump gene expression. Azoles and 5-FC trigger the deacetylation process at the K97 residue of Isw1, initiating the ubiquitin-mediated proteasomal degradation of Isw1 through the E3 ligase Cdc4. The decrease in Isw1 protein level results in the stimulation of drug pump gene expression and the inhibition of 5-FC resistance genes.

The identified ubiquitination sites were classified into three groups based on the domination of Isw1 stability regulation: predominant, moderate, and minor. While Isw1^K441^ was a predominant regulating site of ubiquitination that extensively modulates Isw1 protein degradation and drug resistance, Isw1^K147^, Isw1^K183^, Isw1^K347^, and Isw1^K415^ played a minor role in Isw1 degradation and no roles in drug resistance. Interestingly, Isw1^K113^ played a moderate role in ubiquitination-mediated Isw1 degradation and was identified to have both an acetylation site and a ubiquitination site. A single mutation at Isw1^K113^ had no effect on Isw1 protein levels: K97 acetylation that prevented Cdc4 binding remains intact. The double-mutated Isw1^K97R, K113R^ was protected from degradation, and its protein levels had an increase of 5.6-fold, allowing the *ISW1*^*K97R, K113R*^ strain to be drug resistant. Although it is a challenge to dissect the function of acetylation and ubiquitination at Isw1^K113^, drug treatment represents a deacetylation process for Isw1, and deacetylated Isw1^K113^ is most likely ubiquitinated.

The analysis of clinical isolates revealed a broad regulation phenomenon of Isw1 in drug resistance. Clinical isolates that were more strongly resistant to antifungals produced lower levels of Isw1. In addition, both modifications were identified in tested clinical isolates, demonstrating that Isw1 undergoes acetylation and is subject to the ubiquitin-proteasome pathway. Importantly, overexpressing Isw1 dampened drug resistance in clinical isolates. These discoveries suggested that drug resistance in clinical isolates is modulated by the regulation of Isw1 protein levels.

Taken together, our evidence demonstrates the critical function of Isw1 as a master regulator of multidrug responses in both laboratory and clinical strains. It allows us to decipher the molecular mechanism of the acetylation-Isw1-ubiquitination axis that modulates the expression of drug-resistant genes. These findings underscore the importance of performing thorough evaluations of PPTMs in drug resistance mechanism studies, highlighting a potential strategy for overcoming fungal drug resistance.

## Acknowledgment

We thank Profs. Yongqiang Fan and Ren Sheng for their critical review of the manuscript. This work was supported by the National Key Research and Development Program of China (2022YFC2303000). Funds for this program were also provided by the National Natural Science Foundation of China (31870140 to C.D.) and Liaoning Revitalization Talents Program (XLYC1807001 to C.D.). Research in PW lab was supported by the National Institutes of Health (US) awards AI156254 and AI168867.

## Author contributions

Y.M., T.S., Y.L. and C.D. designed the project. Y.M., Z.L., T.J. and X.G. conducted the experiments. H.L. carried out bioinformatic analyses. C.S., C.L. and S.Y. assisted with molecular biology. G.L. and TB.L. contributed to strain generation. C.D., Y.M. and X.G. participated in data analysis. C.D., P.W. and Y.M. composed the manuscript. All authors reviewed and edited the manuscript.

## Material and Methods

### Ethical statement

All animal experiments were reviewed and ethically approved by the Research Ethics Committees of the National Clinical Research Center for Laboratory Medicine of the First Affiliated Hospital of China Medical University (KT2022284) and were carried out in accordance with the regulations in the Guide for the Care and Use of Laboratory Animals issued by the Ministry of Science and Technology of the People’s Republic of China. Infections with *C. neoformans* were performed via the intranasal route. Four- to six-week-old female Balb/c mice were purchased from Changsheng Biotech (Liaoning, China) and used for survival and fungal burden analyses.

### Strains and growth conditions

Fungal cells **(Table S2)** were routinely grown in YPD medium (1% yeast extract, 2% peptone, 2% dextrose). Biolistic transformation was performed using YPD medium supplemented with 100 μg/ml nourseothricin (WERNER BioAgents), 200 μg/ml neomycin (Inalco) and 200 U/ml hygromycin B (Calbiochem) followed by colony selection. Deacetylases were blocked using 3 μM TSA (MedChemExpress) and 20 mM NAM (Sigma). Drug resistance tests were performed using 16 μg/ml or 20 μg/ml FLC (MedChemExpress), 0.2 μg/ml or 0.3 μg/ml ketoconazole (MedChemExpress), 100 μg/ml or 200 μg/ml 5-FC (MedChemExpress), 0.5 μg/ml or 1.0 μg/ml Amp B (MedChemExpress) or 50 μg/ml or 100 μg/ml 5-FU (MedChemExpress).

### Determination of MIC

Overnight cultures were purified from liquid YPD by washing three times with PBS, then adjusting until OD_600_ was 0.02, then 100 µl of cell suspension was added to each well of a 96-well plate (10,000 cells per well). The well plate was incubated at 30°C for 24 hours, and OD_600_ readings were taken using a Synergy H4 microplate reader (BioTek). The MICs was calculated using XY analyses in GraphPad 6.0.

### Mass spectrometry

Mass spectrometry was performed to analyze Isw1 ubiquitination. The *ISW1-FLAG* complementation strains were subcultured at 30°C with 200 μM MG132 (MedChemExpress) in 50 ml YPD media at 30°C, and cells in the mid-log phase were used. Cell proteins were extracted using lysis buffer (50 mM Tris-HCl, 150 mM NaCl, 0.1% NP-40; pH 7.5) with 1x protease inhibitor cocktail (CWBIO) and 40 mM PMSF. All lysed protein samples were incubated with anti-Flag magnetic beads (Sigma) at 4°C overnight. The beads were washed with TBS buffer (50 mM Tris-HCl, 150 mM NaCl, 1% Triton X-100; pH 7.4) three times, and the bound proteins were extracted into protein loading buffer (125 mM Tris-HCl, 4% SDS, 20% glycerol; pH 7.5) at 95°C for 5 minutes. All protein samples were separated using 8% SDS-PAGE electrophoresis, and the protein gel was stained with Coomassie Brilliant Blue R-250 (BBI Life Sciences) followed by clipping the gel strip.

In-gel tryptic digestion was performed by destaining the gel strips in 50 mM NH_4_HCO_3_ in 50% acetonitrile (v/v) until clear. The gel strips were dehydrated using 100 μl 100% acetonitrile for 5 minutes, rid of liquid, rehydrated in 10 mM dithiothreitol, then incubated at 56°C for 60 minutes. They were again dehydrated in 100% acetonitrile and rid of liquid, then rehydrated with 55 mM iodoacetamide followed by incubation at room temperature in the dark for 45 minutes. Next, they were washed with 50 mM NH_4_HCO_3_, dehydrated with 100% acetonitrile, then rehydrated with 10 ng/μl trypsin and resuspended in 50 mM NH_4_HCO_3_ on ice for 1 hour. After removing excess liquid, they were digested in trypsin at 37°C overnight, then peptides were extracted using 50% acetonitrile/ 5% formic acid followed by 100% acetonitrile.

The peptides were dried completely then resuspended in 2% acetonitrile/ 0.1% formic acid. The tryptic peptides were dissolved in 0.1% formic acid (solvent A) then loaded directly onto a homemade reversed-phase analytical column (15 cm x 75 μm) on an EASY-nLC 1000 UPLC system. They were eluted at 400 nl/min using a gradient mobile phase that increased in solvent B (0.1% formic acid in 98% acetonitrile) from 6% to 23% over 16 minutes, from 23% to 35% over 8 minutes and from 35% to 80% over 3 minutes. Elution continued at 80% for an additional 3 minutes.

The peptides were subjected to an NSI source followed by tandem mass spectrometry (MS/MS) using a Q Exactive™ Plus (Thermo) mass spectrometer coupled to the UPLC. The electrospray voltage applied was 2.0 kV, the m/z scan range was 350 to 1800 for a full scan and intact peptides were detected using an Orbitrap at a resolution of 70,000. Peptides were then selected for MS/MS using an NCE of 28, and the fragments were detected in the Orbitrap at a resolution of 17,500. The data-dependent procedure alternated between one MS scan and 20 MS/MS scans with a 15.0-second dynamic exclusion. The automatic gain control was set at 5E4.

The resulting MS/MS data were processed using Proteome Discoverer 1.3. Spectra were compared against acetylation, ubiquitination or sumoylation databases. Trypsin/P (or other enzymes, if any) was specified as a cleavage enzyme, allowing up to 4 missing cleavages. Mass error was set to 10 ppm for precursor ions and 0.02 Da for fragment ions. Fixed modification was set to cysteine alkylation; variable modifications were set to lysine acetylation, ubiquitination or sumoylation (QEQQTGG and QQQTGG), methionine oxidation and protein N-terminal acetylation. Peptide confidence was set to ‘high,’ and peptide ion score was set to ‘> 20.’

### Strain generation

*C. neoformans* mutants were generated using the H99 strain and biolistic transformation (Toffaletti et al., 1993). The neomycin or nourseothricin resistance marker was amplified using primers M13F and M13R **(Table S3)**. The upstream and downstream DNA sequences of the target gene and the selective marker sequence were joined using overlapping PCR. The resulting PCR fragments were purified, concentrated and transformed into the H99 strain using biolistic transformation. Transformants were selected on YPD agar supplemented with either neomycin or nourseothricin. Correct integration and the loss of target DNA sequences were confirmed using diagnostic PCR.

The *ISW1* gene (*CNAG_05552*) was disrupted by the homologous replacement of its open reading frame (ORF) with a piece of DNA containing a dominant drug resistance gene marker as described henceforth. In the first round of PCR, the primer pairs 3534/3535 and 3536/3537 were used to amplify the 5’ and 3’ flanking regions, respectively, of the *ISW1* gene. The gel-extracted DNA fragments from the first round of PCR were used as templates, and the *isw1Δ::NEO* construct was amplified using the 3534/3537 primer pair. The H99 strain of *C. neoformans* was biolistically transformed with the deletion allele. To identify the desired *isw1Δ* mutant, diagnostic PCR was performed using the 3534/3537 primer pair, and real-time PCR followed, using primers 3557/3558. The same method was used to construct knockout strains of the E3 ligase-related genes in the *ISW1*^*K97R*^ strain. Briefly, the upstream or downstream genomic DNA sequences of the target genes were amplified using the primers listed in the Supplementary Data primer table.

The *ISW1-FLAG* complementation strains were generated as described henceforth. The downstream genomic DNA sequence of *ISW1* was amplified using primers 3735/3736 and cloned between the restriction sites *Sac*II and *Sac*I into the pFlag-NAT plasmid (a plasmid containing the nourseothricin resistance marker and the Flag tag). The complete ORF of the target gene (including the promoter sequence) was amplified using primers 3733/3734 and cloned between the restriction sites *Hind*III and *EcoR*I into pFlag-NAT-dw. The cassette amplified from the final pFlag-NAT by primer 3898-MY/3899-MY was biolistically transformed into the *isw1Δ::NEO* strain. Diagnostic PCR was performed using the 3733/3734 primer pair. Real-time PCR analysis was performed using a gene-specific probe amplified using the 3557/4421-MY primer pair. Western blot analysis of Isw1 was performed using anti-Flag mouse monoclonal antibodies.

The R mutation was formed using site-directed mutagenesis approaches. The K89 codon was first mutated using primers 3733/3866-MY and 3891/3734, then the K89R construct was amplified using the 3733/3734 primer pair and cloned between the restriction sites *Hind*III and *Eco*RI into pFlag-NAT-dw. The K97 codon was mutated using primers 3733/3787 and 3862-MY/3734, whereas the K113 codon was mutated using primers 3733/3836 and 3864-MY/3734. The resulting plasmids contained single-point mutants (K89R, K97R or K113R), double-site mutants (K89R, K97R; K89R, K113R or K97R, K113R) and triple-site mutant (K89R, K97R, K113R). The Q mutation was formed using the same procedures except that the K89Q construct was formed using the primer pair 3733/3866-MY and 3892/3734, K97Q was formed using the primer pair 3733/3787 and 3863-MY/3734 and K113Q was formed using the primer pair 3733/3836 and 3865-MY/3734. The resulting plasmid contained single-point mutants (K89Q, K97Q or K113Q), double-site mutants (K89Q, K97Q; K89Q, K113Q or K97Q, K113Q) and triple-site mutant (K89Q, K97Q, K113Q). All mutant plasmids were further confirmed using DNA sequencing. The cassette amplified from the final pFlag-NAT by primer 3898-MY/3899-MY was biolistically transformed into the *isw1Δ::NEO* strain. Mutant strains were confirmed using DNA sequencing, diagnostic PCR, qRT-PCR and immunoblotting.

Ubiquitination mutants were also formed. The mutant K147R was formed using primer pair 3733/4445-MY and 4444-MY/3734, whereas K183R was amplified using the primer pair 3733/4447-MY and 4446-MY/3734 and K297R was mutated using primers 4448-MY/4449-MY. Similarly, K347R was mutated using primers 4450-MY/4451-MY, K415R was mutated using primers 3807-MY/3809-MY and K441R was mutated using primers 4452-MY/4453-MY. The resulting plasmids were used to generate the R mutant plasmid using the TaKaRaMutanBEST Kit (Takara). All strains were validated using the methods described earlier.

To demonstrate the direct protein interaction between Isw1 and Cdc4, the downstream genomic DNA sequence of *CDC4* was amplified using primers ZR34/ZR51 and cloned between the restriction sites *Spe*I and *Sac*I into the pHA-HYG plasmid (a plasmid containing the hygromycin B resistance marker and the HA tag). Then, the last 1000 bp of the *CDC4* ORF was amplified using primers ZR48 and ZR33, and the resulting fragment was cloned into the above plasmid between *Cla*I and *Sma*I. The cassette amplified from the final plasmid by primers ZR48 and ZR51 was biolistically transformed into H99, the *ISW1*^*WT*^, *ISW1*^*K97Q*^ and *ISW1*^*K97R*^ strains. Diagnostic PCR was performed using the ZR48/ZR51 primer pair. Immunoblotting analysis of Cdc4 was performed using anti-HA (C29F4) rabbit mAb.

To detect the protein expression levels of Isw1 in clinical strains, the wild-type plasmid with pFlag-NAT was used as a template in PCR using primers 3557 and 3537. The resulting PCR products were transformed into seven clinical strains. The *ISW1*^*K97Q*^ overexpression strains were generated as described henceforth. A safe-haven site was applied to perform plasmid integration (Arras et al., 2015). The 3’ flanking region of the safe haven was amplified using 4470-Y/4471-MY, then cloned into pFlag-NAT between the *Sac*II and *Sac*I sites. The 5’ flanking region of the safe haven was amplified using primer 4800-MY/4467-MY, the *TEF1* promoter was amplified using primer 4468-MY/2342 and the *ISW1*^*K97Q*^ coding sequence was amplified using 4807-MY and 3736. The *ISW1*^*K97Q*^ construct was amplified using the 4800-MY/3736 primer pair, and the three gel-extracted DNA fragments from the first and second rounds of PCR were used as templates, then cloned into pFlag-NAT-dw between the *Hind*III and *EcoR*I sites. The *ISW1*^*K97Q*^ overexpression cassette was amplified using the 4800-MY/4471-MY primer pair, and the product was transformed into clinical strains.

### Animal infection

Mice were anaesthetized and inoculated intranasally with 10^5^ yeast cells suspended in 50 μl PBS buffer. Infected mice were weighed 12 days after infection and then monitored twice daily for morbidity. Mice were sacrificed at the endpoint of the experiment. All animal experimentation was carried out under the approved protocol (please see Ethical statement).

### Transcriptome and qRT-PCR analyses

To analyze drug resistance, the wild-type H99 and *isw1Δ* mutant strains were either untreated or were treated with 10 μg/ml FLC (Sigma) in 50 ml YPD media at 30°C until cell densities reached the exponential phase (approximately 6 to 7 hours). Cells were then washed three times with ice-cold PBS and placed in a tank of liquid nitrogen. Total RNA was isolated using TRIzol reagent (Thermo Fisher Scientific), and 3 μg of the product was processed using the TruSeq RNA Sample Preparation Kit (Illumina). Purification of mRNA was performed using polyT oligo-attached magnetic beads. Fragmentation of mRNA was performed using an Illumina proprietary fragmentation buffer. First-strand cDNA was synthesized using random hexamer primers and SuperScript II. Subsequently, second-strand cDNA was synthesized using RNase H and DNA polymerase I. The 3′ end of the cDNA sequence was adenylated, then cDNA sequences of 200 bp were purified using the AMPure XP system (Beckman Coulter) and enriched using an Illumina PCR Primer Cocktail in a 15-cycle PCR. The resulting PCR products were then purified, and integrity was confirmed using an Agilent High Sensitivity DNA assay on a Bioanalyzer 2100 (Agilent). The sequencing library was then sequenced using a Hiseq platform (Illumina) by Shanghai Personal Biotechnology Cp. Ltd. Alignments were checked against the *Cryptococcus_neoformans*_var._*grubii_*H99 reference genome and gene annotation set retrieved from Ensemble. Differentially expressed genes were detected using the Bioconductor package DESeq2 version 1.22.2. Genes with adjusted *p* values <0.05 and changes greater or less than 1.5-fold those of the control strain were considered to be significantly induced or repressed, respectively.

To verify the gene changes screened by transcriptomics, H99 and *isw1Δ* mutant strains were either untreated or were treated with 40 μg/ml FLC (MedChemExpress) in 10 ml YPD media at 30°C, and cell densities were monitored until OD_600_ reached 1.0. Both the H99 and *isw1Δ* mutant strains were either untreated or treated with 400 μg/ml 5-FC (MedChemExpress) in 10 ml YPD media at 30°C for 1 hour. Cells were harvested at 3000 *rpm* for 3 minutes at 4°C, then washed twice with ice-cold PBS. Total RNA was isolated using a total RNA kit I (Omega), and cDNA was synthesized using a reverse transcript all-in-one mix (Mona). Primers for amplifying target genes can be found in the primer table. Data were acquired using a CFX96 real-time system (Bio-Rad) using actin expression as a normalization control. The *ΔΔCt* method was used to calculate differences in expression.

### Co-immunoprecipitation and immunoblotting assays

Overnight cultures of *C. neoformans* strains were diluted in fresh YPD media and incubated at the indicated temperature to the mid-log phase (OD_600_=0.8). Protein immunoprecipitation or co-immunoprecipitation was performed as described elsewhere (Li YJ et al., 2017). Briefly, cell proteins were extracted using lysis buffer (50 mM Tris-HCl, 150 mM NaCl, 0.1% NP-40; pH 7.5) with 1X protease inhibitor cocktail (CWBIO) and 40 mM PMSF. Aliquots of protein extracts were retained as input samples. Samples of the lysed protein were incubated with anti-Flag magnetic beads (MedChemExpress) at 4°C overnight, then the beads were washed three times using TBS buffer, and the bound proteins were extracted into protein loading buffer at 95°C for 5 minutes. Protein samples were separated using 8% SDS-PAGE electrophoresis, transferred onto nitrocellulose membranes, and blocked using 5% milk. Immunoblotting or co-immunoprecipitation assays were performed using anti-Flag mouse monoclonal antibodies (1:5000 dilution; Transgene), anti-HA (C29F4) rabbit mAb (1:5000 dilution; Cell Signaling Technology), anti-Histone H3 (D1H2) XP® Rabbit mAb (1:5000 dilution; Cell Signaling Technology), goat anti-mouse IgG (H+L) HRP secondary antibodies, and goat anti-rabbit IgG (H+L) HRP secondary antibodies (1:5000 dilution; Thermo Fisher Scientific), and monoclonal and polyclonal Kac (1:2500; PTM Bio). The signal was captured using a ChemiDoc XRS+ (Bio-Rad).

### Statistical analysis

All statistical analyses were performed using GraphPad Prism software (GraphPad 6.0). Two-tailed unpaired *t* tests were used in two-sample comparisons. Statistical analyses for two or more groups were performed using one-way or two-way ANOVA. Significant changes were recognized when *p* < 0.05. All experiments were performed using at least three biological replicates to ensure reproducibility.

### Materials and Data availability

The raw Isw1 proteome modification mass spectrometric data have been deposited to the Proteome Xchange (https://www.ebi.ac.uk/pride) with identifier PXD037150 (username, password: flU9d0tA). The transcriptome (RNA-seq) is deposited in NCBI’s Gene Expression Omnibus (GEO) (https://www.ncbi.nlm.nih.gov/geo/) and can be accessed through GEO Series accession ID GEO:GSE217187. Any other data necessary to support the conclusions of this study are available in the supplementary data files and source data or are available from the authors upon request.

### Detection of drug content

To analyze drug resistance, the wild-type H99 and *isw1Δ* mutant strains were treated with 40 μg/ml FLC in 50 ml YPD media at 30°C until the cell densities reached the exponential phase (approximately 5 hours), then the cells were washed once with PBS. An appropriate amount of uniform sample was weighed, 0.2 ml 50% acetic acid solution was added, ultrasonic extraction was carried out, and the resultant was passed through a 0.22 μm microporous filter membrane. High-performance liquid chromatography was performed using an injection volume of and a constant mobile phase flow rate of 1.0 ml/min. An Agilent C18 (4.6 mm x 250 mm x 5 μm) column was used, held at 35°C, on a Thermo U3000 HPLC; the detector was a DAD. When the drug was FLC, the mobile phase was acetonitrile:water:acetic acid (25:75:0.2), the detector wavelength was 261 nm, the run time was 15 minutes, and the standard curve was *Y*=0.0147*X*-0.0109 (*r*^*2*^=0.9999).

The drug content was determined as:

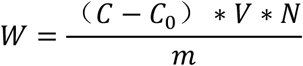

*W*——Drug content, mg/kg

*C*——Concentration of the drug in the cell, mg/L

*C*_*0*_——Concentration of the drug in the blank control, mg/L

*V*——Volume, ml

*N*——Diluted

`*m*——cell mass, g

## Figures and legends

**Figure 1-figure supplement 1.**
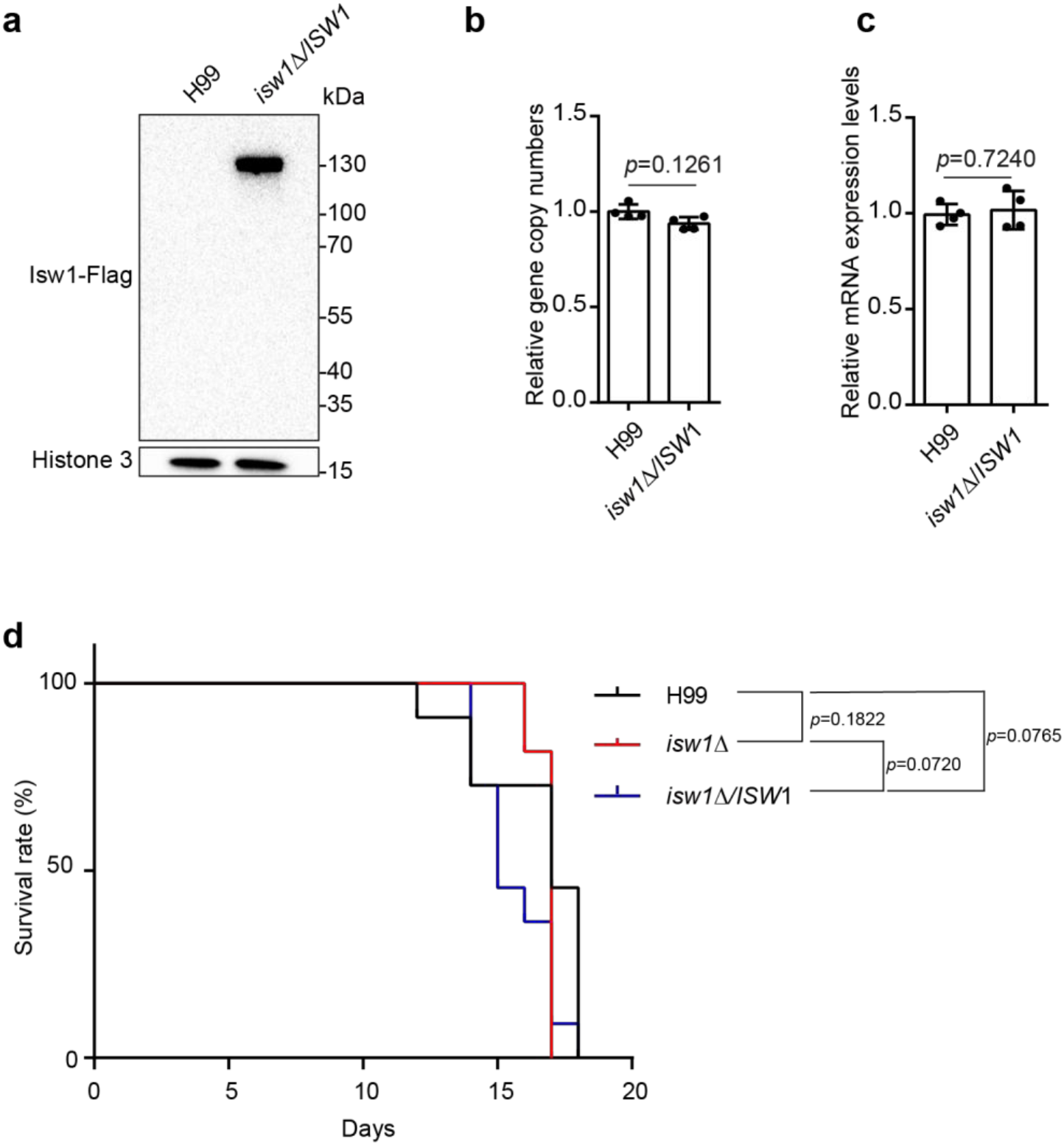
*ISW1* is not required for fungal virulence. a. Immunoblotting analysis of the *ISW1* complementation strain. The complementation strain was constructed, and an immunoblotting assay was performed to confirm the expression of the Isw1-Flag protein. b. Genomic copy number assays. Genomic DNA was isolated from the wild-type and complementation strains, then qRT-PCR was performed on each to confirm the gene copy number of *ISW1*. Oligos of actin were used as a control. Four independent assays were performed and quantified. Two-tailed unpaired *t-*tests were used. Data are expressed as mean ± SD. c. Analyses of the *ISW1* complementation strain using qRT-PCR. Samples of RNA were isolated from the wild-type and complementation strains, then qRT-PCR was performed on each to confirm the gene expression of *ISW1*. Oligos of actin were used as a control. Four independent assays were performed and quantified. Two-tailed unpaired *t-*tests were used. Data are expressed as mean ± SD. d. Animal survival analysis and the Kaplan-Meier survival curves of wild-type and *isw1Δ*. Significance was determined using a log-rank (Mantel-Cox) test.

**Figure 1-figure supplement 2.**
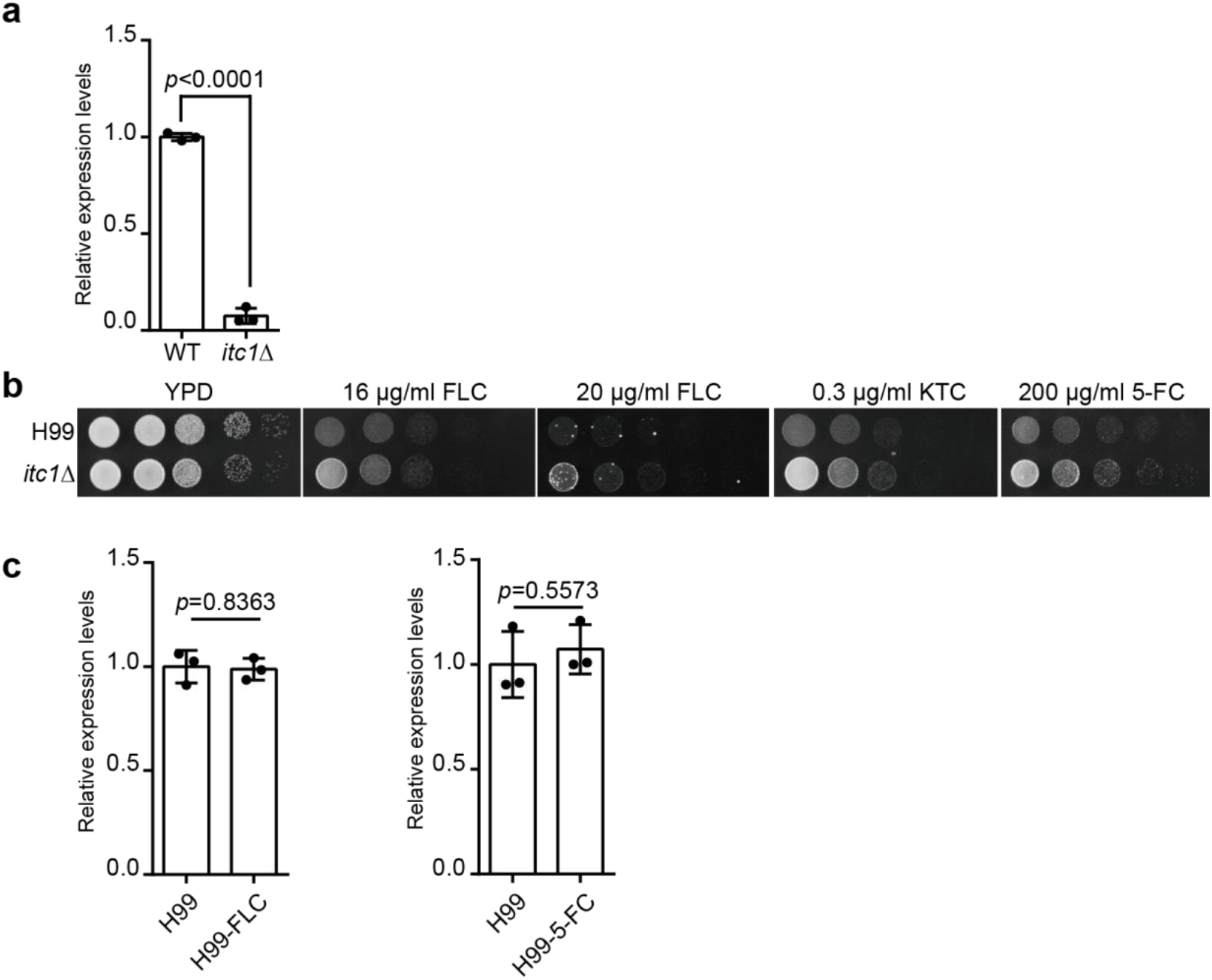
*itc1Δ* is resistant to azoles and 5-fluorocytosine. a. Analyses of *ITC1* in the *itc1Δ* strain using qRT-PCR. Samples of RNA were isolated from the wild-type and *itc1Δ* strains, then qRT-PCR was performed on each. Two-tailed unpaired *t-*tests were used. Data are expressed as mean ± SD. b. Spotting assays of *itc1Δ*. The wild-type and *itc1Δ* strains were separately spotted onto YPD agar either supplemented with an antifungal agent or left blank. c. Analyses of *ISW1* gene expression in response to FLC and 5-FC. Samples of RNA were isolated from the wild-type strain, then qRT-PCR was performed on each. Two-tailed unpaired *t-*tests were used. Data are expressed as mean ± SD.

**Figure 3-figure supplement 1.**
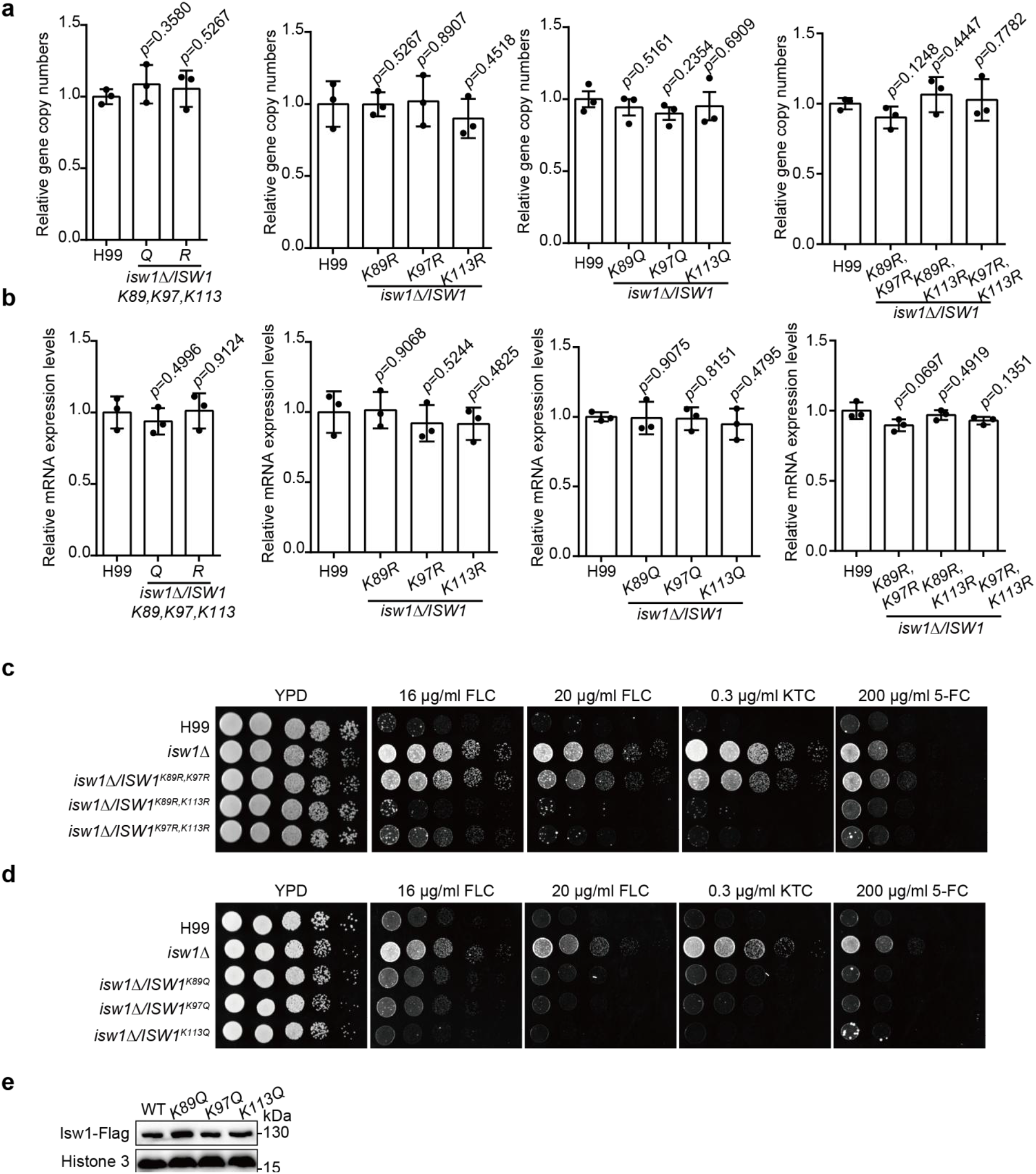
Screening important acetylation sites of Isw1. a. Genomic copy number assays. Genomic DNA was isolated from the indicated *ISW1* mutants, then qRT-PCR was performed on each to confirm the gene copy number of *ISW1*. Oligos of actin were used as a control. Three independent assays were performed and quantified. Two-tailed unpaired *t-*tests were used. Data are expressed as mean ± SD. b. Analyses of *ISW1* mutants using qRT-PCR. Samples of RNA were isolated from the wild-type and indicated mutants, then qRT-PCR was performed on each to confirm the gene expressions of wild-type and mutated *ISW1*. Oligos of actin were used as a control. Three independent assays were performed and quantified. Two-tailed unpaired *t-*tests were used. Data are expressed as mean ± SD. c. Spotting assays of *ISW1* double-R mutants. The wild-type and indicated *ISW1* mutants were spotted onto YPD agar either supplemented with an antifungal agent or left blank. d. Spotting assays of *ISW1* single-Q mutants. The wild-type and indicated *ISW1* mutants were spotted onto YPD agar either supplemented with an antifungal agent or left blank. e. Immunoblotting analyses of *ISW1* single-Q mutants. Protein samples were isolated from the indicated strains, and immunoblotting assays were performed.

**Figure 4-figure supplement 1.**
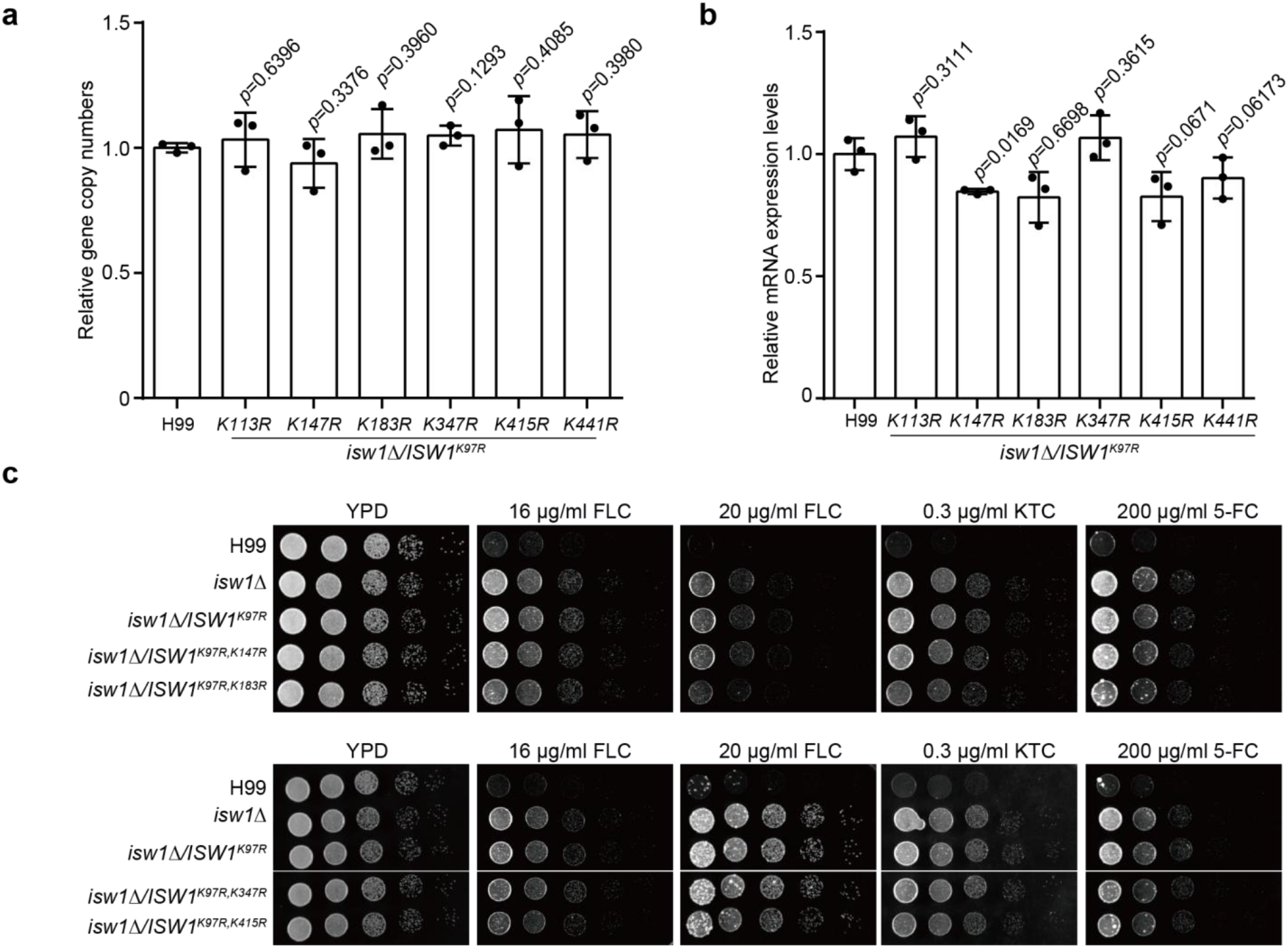
Screening important ubiquitination sites of Isw1. a. Genomic copy number assays. Genomic DNA was isolated from indicated *ISW1* mutants, then qRT-PCR was performed on each to confirm the gene copy number of *ISW1*. Oligos of actin were used as a control. Three independent assays were performed and quantified. Two-tailed unpaired *t-*tests were used. Data are expressed as mean ± SD. b. Analyses of *ISW1* ubiquitination mutants using qRT-PCR. Samples of RNA were isolated from wild-type and indicated mutants, and qRT-PCR was performed on each to confirm the expressions of wild-type and mutated *ISW1* genes. Oligos of actin were used as a control. Three independent assays were performed and quantified. Two-tailed unpaired *t-* tests were used. Data are expressed as mean ± SD. c. Spotting assays of *ISW1* ubiquitination site mutants. The wild-type and indicated *ISW1* mutants were spotted onto YPD agar either supplemented with an antifungal agent or left blank. Plates were incubated at 30°C for 3 days.

**Figure 5-figure supplement 1.**
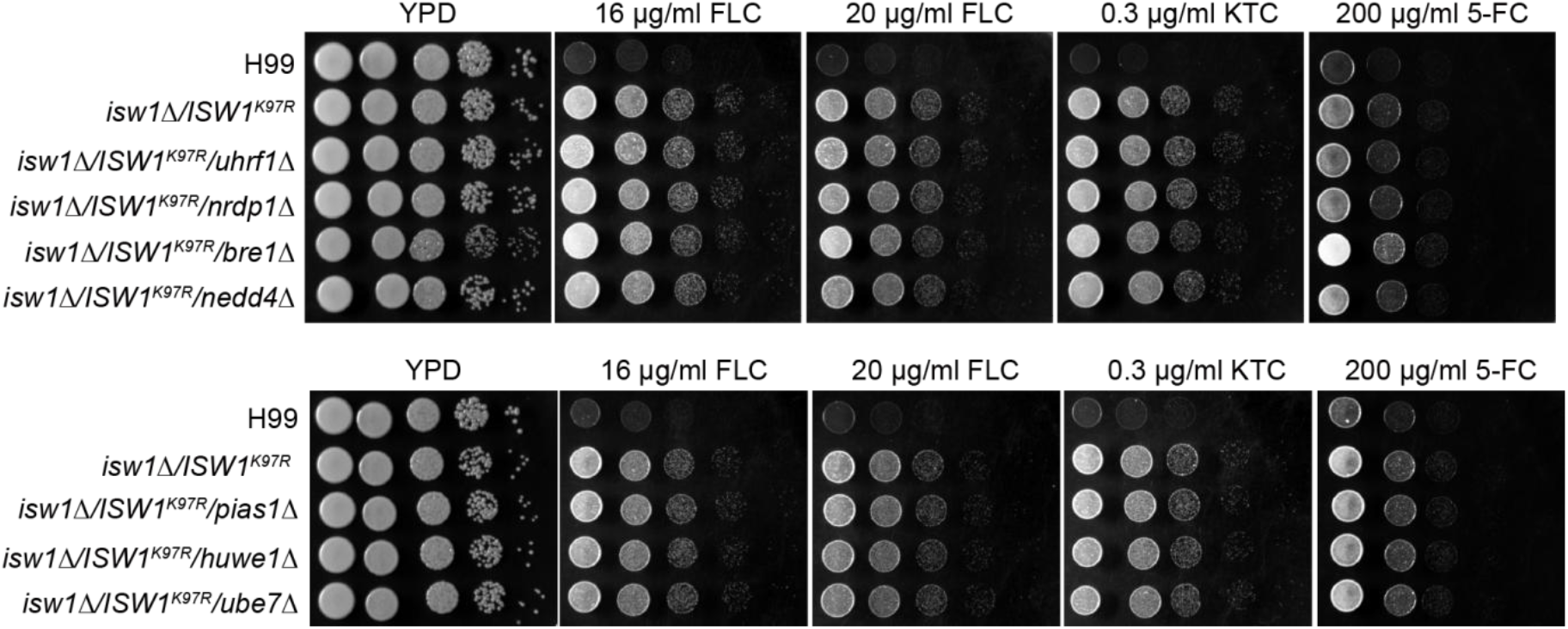
Identification of the E3 ligase for Isw1. Spotting assays of E3 ligase knockout strains. Wild-type and indicated E3 ligase knockout strains were spotted onto YPD agar either supplemented with an antifungal agent or left blank. Plates were incubated at 30°C for 3 days.

**Table S1. DEGs (differentially expressed genes) in *isw1/****Δ* **cells treated with FLC**.

**Table S2. Strains used in this study**.

**Table S3. Primers used in this study**.

